# Salient Experiences are Represented by Unique Transcriptional Signatures in the Brain

**DOI:** 10.1101/192278

**Authors:** Diptendu Mukherjee, Bogna Marta Ignatowska-Jankowska, Eyal Itskovits, Ben Jerry Gonzales, Hagit Turm, Liz Izakson, Doron Haritan, Noa Bleistein, Chen Cohen, Ido Amit, Tal Shay, Brad Grueter, Alon Zaslaver, Ami Citri

## Abstract

Inducible transcription is essential for consolidation of salient experiences into long-term memory. However, the question of whether inducible transcription relays information representing the identity of the experience being encoded, has not been explored. To this end, we have analyzed transcription across multiple brain regions, induced by a variety of rewarding and aversive experiences. Our results define robust transcriptional signatures uniquely characterizing individual salient experiences. A subset of these induced transcriptional markers suffice for near-perfect decoding of the identity of recent experiences at the level of individual mice. Furthermore, experiences with shared attributes display commonalities in their transcriptional representation, exemplified in the representation of valence, habituation and reinforcement. Taken together, our results demonstrate the existence of a neural transcriptional code that represents the encoding of experiences in the mouse brain. This code is comprised of distinct transcriptional signatures that correlate to the affective attributes of the experiences that are being encoded.

## Introduction

Neuronal plasticity enables cognitive and behavioral flexibility underlying the development of adaptive behaviors^1,2^. This neuroplasticity, induced by salient experiences, has been shown to depend on the induction of temporally-defined waves of transcription^1–5^. The earliest of these waves consists of the expression of immediate-early genes (IEGs). IEGs have been conventionally treated as molecular markers for labelling neuronal populations that undergo plastic changes, underlying the formation of long-term memory^6,7^. However, recent literature indicates a much more significant contribution of IEGs in synaptic plasticity and memory formation^8,9^. It has been proposed that IEG transcription could represent the molecular signatures of long-term plastic changes underlying the formation of memory^1^. This implies that IEG expression induced by an experience could represent a neural transcriptional code for long-term storage of information that is specific to an experience. The existence of a neural code embedded in transcription implies that it should be possible to decode the identity of recent experiences, and potentially derive information regarding the nature of the experience, from its transcriptional representation^10^. This proposition forms the basis of our investigation.

To address the existence of a neural transcriptional code, we performed detailed analysis of IEG transcription for 14 different experiences (see methods for details), induced by cocaine (acute, repeated and challenge), volitional sucrose drinking (acute and repeated), reinstatement of feeding following food deprivation, lithium chloride administration (LiCl; acute and repeated), saline (acute injection without habituation, acute injection after habituation and following repeated administrations), acute administration of a mild foot shock, and exposure to a novel chamber with no foot shock. The experiences analyzed were selected to enable identification of the transcriptional representations of affective attributes characterizing an experience, such as valence and salience^11,12^. Analysis of repeated exposure to experiences also enabled identification of common attributes in the transcriptional representation of habituation and positive reinforcement.

Assuming that encoding of complex behaviors involves the coordinated transcriptional activation of multiple brain regions, we analyzed transcription across multiple structures of the reward circuitry^13^. The brain structures that were analyzed include limbic cortex (LCtx; including medial prefrontal cortex and anterior cingulate cortex), nucleus accumbens (NAc), dorsal striatum (DS), amygdala (Amy), lateral hypothalamus (LH), dorsal hippocampus (Hipp) and ventral tegmental area (VTA).

Our results demonstrate that the transcriptional representations of each experience are robust, reliable and consistent, enabling the decoding of the recent salient experience of mice with high levels of accuracy from a minimal transcriptional signature. We identify transcriptional correlates for affective attributes of experience, prominently demonstrated in the encoding of valence. Moreover, we report opposing patterns of transcriptional modulation underlying the development of habituation to experiences of neutral or negative valence, in comparison to reinforcement for repeated rewarding experiences. We finally discuss the potential implications of the identification of a neural transcriptional code for salient experiences.

## Results

### Unique transcriptional signatures represent the history of cocaine experience

Drugs of abuse are known to hijack endogenous mechanisms of neural plasticity, inducing long-lasting modifications of neural circuits in the mesolimbic dopamine system^14^ through transcription-dependent mechanisms^15^. Cocaine sensitization is one of the most widely applied paradigms for studying mechanisms of neural plasticity, due to the robustness of the behavioral model and the detailed insight provided into the underlying mechanisms^3,15–20^. We initiated our study with an investigation of the gene expression programs induced during the development of behavioral sensitization to cocaine.

Using the cocaine sensitization paradigm, we studied the transcriptional programs induced in mice following acute exposure to cocaine (intraperitoneal (i.p.) injections, 20 mg/kg), or repeated exposure to the drug, characterized by robust locomotor sensitization. The response to re-exposure to cocaine after a period of abstinence from repeated drug exposures (‘cocaine challenge’), is a classic metric of the potent experience-dependent maladaptive plasticity induced by cocaine exposure^15^ (***Figure 1A,B***). We analyzed transcriptional dynamics at 0, 1, 2, 4 hrs following each of these cocaine experiences across 7 brain structures (LCtx, NAc, DS, Amy, LH, Hipp and VTA, applying a comprehensive set of qPCR probes developed against putative IEGs (**Table S1**). We observed that the transcriptional representation of distinct cocaine experiences (acute, repeated, challenge) was characterized by the robust induction of a handful of genes across brain structures (***Figure 1C, Figure 1 - Figure Supplement 1***). A subset of these genes (*Arc*, *Egr2*, *Egr4*, *Fos* and *Fosb*) displayed consistently high induction and low variance, as well as clear temporal dynamics of induction, with expression peaking at 1 hour following cocaine experience (***Figure 1 - Figure Supplement 2***). Importantly, these five genes were robustly induced across all salient experiences investigated in this work, and were therefore chosen as the representative markers of the recent experience of individual mice for the rest of the study.

**Figure 1.**
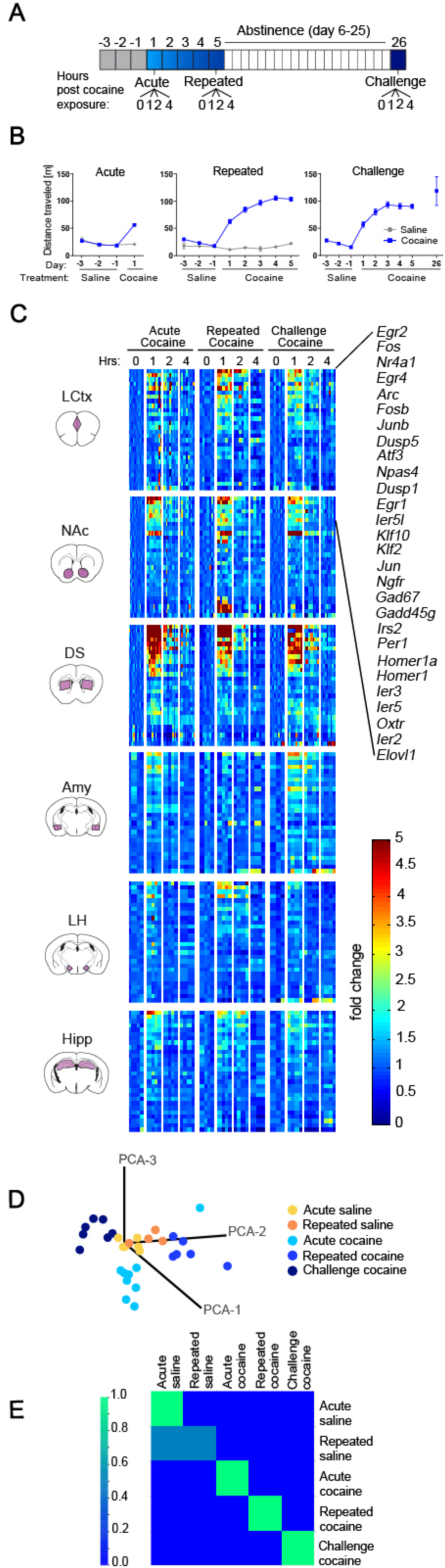
Specific transcriptional signatures encode cocaine experiences. **(A)** Schematic of experimental paradigm. Mice were exposed to cocaine (i.p., 20 mg/kg) or saline, either acutely, repeatedly or re-exposed after abstinence (challenge), with transcriptional dynamics studied at 0, 1, 2 or 4 hrs. **(B)** Locomotor activity of mice following acute, repeated or challenge cocaine experiences (compared to saline). Sample size: acute saline n = 6; acute cocaine n = 30; repeated saline n = 4; repeated cocaine n = 22; challenge cocaine n = 19 mice. Results indicate mean ± s.e.m. **(C)** Expression matrix of IEG induction dynamics following cocaine experiences. Individual animals are represented in columns sorted according to time points of cocaine experiences [sample numbers per time point - LCtx: limbic cortex (n = 5-11), NAc: nucleus accumbens (n = 5-12), DS: dorsal striatum (n = 5-12), Amy: amygdala (n = 3-4), LH: lateral hypothalamus (n = 2-4), Hipp: hippocampus (n = 2-4)]. Fold induction is graded from blue (low) to red (high). Genes represented were induced at least 1.5-fold over control in any one of the brain regions studied. **(D)** Linear projection of the three principal components after dimension reduction of transcriptional induction of *Arc*, *Egr2*, *Egr4*, *Fos* and *Fosb* in the LCtx, NAc and DS segregates cocaine-treated mice into discrete clusters associated with recent experiences. Each dot represents an individual animal, color-coded according to the identity of its recent experience. **(E)** Confusion matrix representing the classification accuracy of decoding the recent experience (acute / chronic / challenge cocaine vs saline) of individual mice based on expression of *Arc*, *Egr2*, *Egr4*, *Fos* and *Fosb* induction in the LCtx, NAc and DS. Accuracy is scaled from blue to green, with bright green corresponding to 100% accuracy (n = 29 mice).

As a preliminary test of the hypothesis that experiences can be decoded from patterns of induced transcription, we performed supervised classification of animals that experienced distinct saline and cocaine experiences. Mice were classified based on induction of the five most prominently induced genes (*Arc*, *Egr2*, *Egr4*, *Fos* and *Fosb*) across three brain structures (LCtx, NAc and DS). Each gene-structure combination was identified as a “feature” and classification was performed using the k-Nearest Neighbor classifier. Linear projection following dimension reduction of these 15 features into three principal components, resulted in unique clustering of the acute, repeated and challenge experiences of cocaine (***Figure 1D*)**. More importantly, the information contained in these 15 features, allowed precise allocation/classification of individual animals based on the identity of the recent cocaine experiences (***Figure 1E*).** However, while saline experiences could easily be segregated from cocaine experiences, using this approach and the given set of features, we could only partially segregate between the neutral experiences induced by exposure to acute or repeated saline. We attribute this inability to accurately decode the saline experiences to the effect of habituation of the mice to a neutral stimulus, rendering the experience of repeated saline injections non-salient and therefore not represented by inducible transcription (***Figure 1 - Figure Supplement 3*, 4**).

Taken together, these results suggest that transcriptional signatures comprising of a minimal subset of marker genes reliably represent the comprehensive gene expression programs induced by experience, and suffice to decode the recent salient experience at the resolution of individual mice.

### Distinct experiences are represented by unique transcriptional signatures

To test this fundamental concept, we expanded our study, including experiences that were characterized by negative valence such as LiCl and foot shock, along with naturalistic volitional experiences of positive valence – sucrose consumption, and reinstatement of feeding following food withdrawal. We represent the experience-specific IEG expression patterns induced 1 hour following an experience across multiple brain nuclei using radar plots (***Figure 2***). This representation provides a birds-eye view of the transcriptional landscape, and enables immediate identification of the major brain regions and transcripts recruited by each experience. Four genes (*Arc, Egr2, Egr4* and *Fos*) were shown for simplicity of presentation; for the data from individual mice, see ***Figure 2 - Figure Supplement 1***.

**Figure 2.**
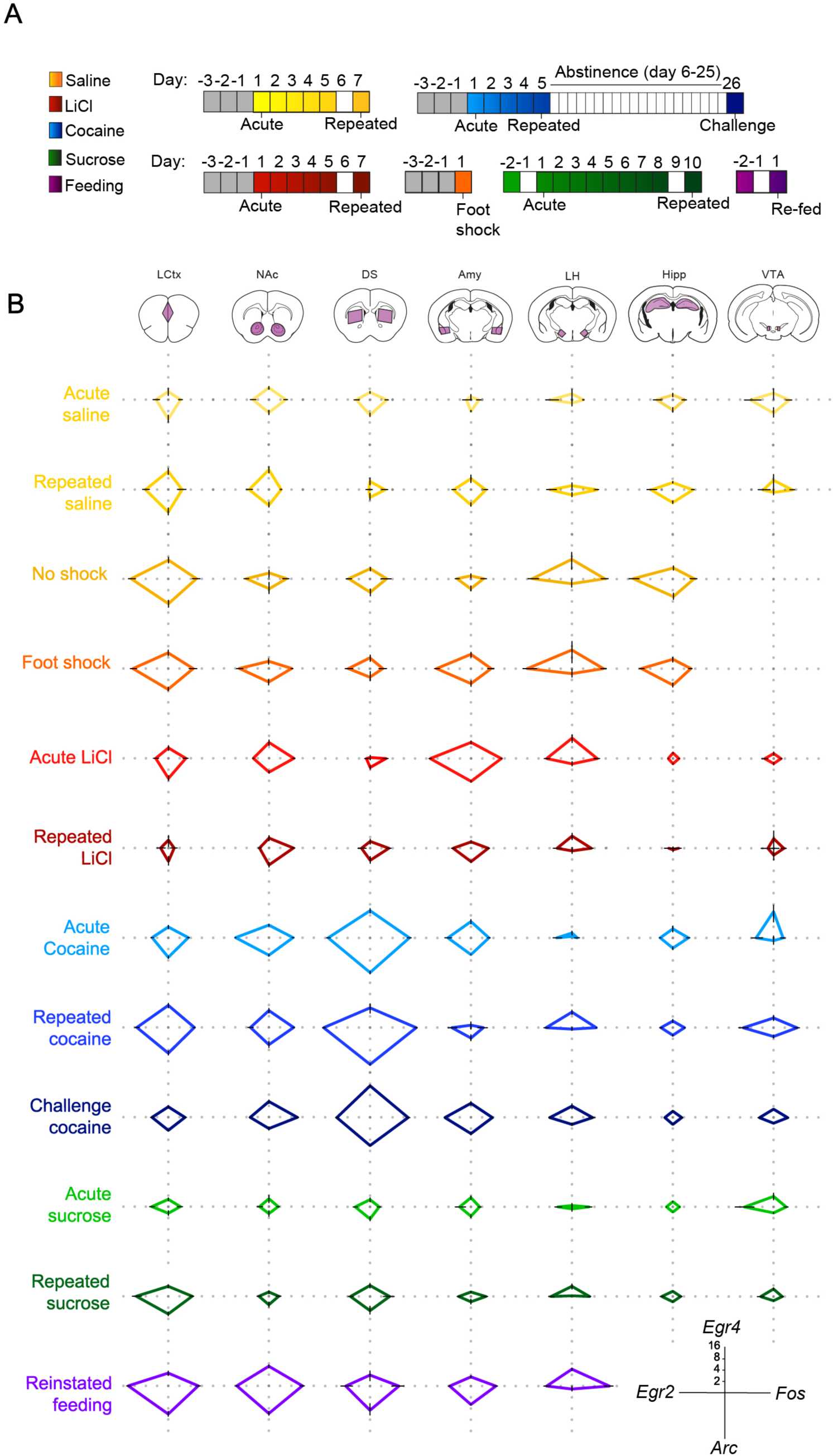
Salient experiences are represented by unique transcriptional signatures. **(A)** Schematic of experimental paradigms. Experiences analyzed include saline (acute & repeated); foot shock (acute shock and no-shock controls exposed to the same environment); LiCl (acute and repeated); cocaine (acute, repeated and challenge following abstinence); sucrose (acute and repeated) and reinstatement of feeding (following 18 hrs of deprivation). **(B)** Radar plots representing the transcriptional induction of *Arc*, *Egr2, Egr4* and *Fos* across seven brain structures 1 hour after the different experiences [LCtx: limbic cortex (n = 4-14), NAc: nucleus accumbens (n = 4-14), DS: dorsal striatum (n = 4-14), Amy: amygdala (n = 4-9), LH: lateral hypothalamus (n = 3-9), Hipp: hippocampus (n = 4-9); VTA: ventral tegmental area (n = 2-8)]. Results are shown in log_2_ scale as mean ± s.e.m of induction over baseline control.

This presentation further highlights the unique nature of the transcriptional code characterizing each experience, and the dynamic changes in IEG induction following repeated exposures to an experience. Furthermore, commonalities in the transcriptional representation of experiences with shared affective attributes were also visually accessible in this presentation.

### The transcriptional representation of negative valence

Investigating the transcriptional representation of negative valence, we focused on the aversive experiences induced either pharmacologically by LiCl administration, or by acute administration of mild foot shock. While animals that received LiCl display malaise, nausea and reduced locomotion^21^; administration of mild foot shock induces acute pain, fear and consequent freezing^22^. To facilitate a direct comparison of the transcriptional patterns induced by experiences of opposing valence, LiCl administration was performed using the same experimental setup as cocaine sensitization. The experiences of cocaine and acute LiCl exposure drove robust induction of a small but similar subset of IEGs, which most prominently included *Arc, Egr2, Egr4, Fos* (***Figure 2 - Figure Supplement 2***). However, in comparison to the cocaine experiences, where transcriptional induction was observed to be pronounced in the LCtx, NAc, DS, and VTA, the LiCl experiences displayed induction predominantly in the Amy (***Figure 2***).

We next compared the experience of LiCl shared with a different aversive experience, induced by foot shock. It is worth noting that while LiCl and foot shock are both characterized by negative valence, they are otherwise distinct in multiple aspects – LiCl causes prolonged visceral discomfort, while foot shock associates acute peripheral pain with a novel environment (a metal grid located in a small enclosure). Interestingly, exposure to the context in which the foot shock was performed, alone, induced robust IEG transcription in a number of limbic structures, (***Figure 2***). Mice that received a foot shock within this context displayed an overall indistinguishable pattern of transcriptional induction compared to their no-shock controls, with the sole distinction being a robust induction of transcription (predominantly of *Egr2* and *Egr4*) in the Amy (***Figure 2*, *Figure 2 - Figure Supplement 3***). This result provides correlative evidence for transcription-dependent encoding of negative valence in the amygdala, introduced by the addition of a single variable (foot shock) to a complex experience of transfer to a novel environment.

Taken together, these results support the notion that recent salient experiences are characterized by robust induction of specialized transcription programs in relevant brain structures. From a broader perspective, our results demonstrate that experiences of opposing valence induce distinct IEG expression patterns in different brain structures (***Figure 2***). The representation of rewarding experiences are characterized by robust transcriptional induction in the LCtx, NAc, DS, and VTA, while the representations of aversive experiences are dominated by transcriptional induction in the Amy.

### Transcriptional representation of habituation and reinforcement

Repeated experience can either cause habituation or reinforcement, depending on the valence of the experience – habituation is observed to experiences of neutral and negative valence, while reinforcement is expressed in response to repeated experiences of positive valence. We therefore expected to observe a reflection of these distinct behavioral trajectories in the transcriptional representation of repeated experiences of positive or negative valence.

Indeed, further analysis identified opposing trajectories of IEG induction following repetition of aversive or rewarding experiences. In case of the aversive LiCl experiences, following repeated exposure, we observed a significantly diminished transcriptional representation in the Amy to levels similar to those observed following repeated saline experience [interaction of treatment (LiCl vs saline) and time (acute vs repeated); *Egr2*: *F*_(1,18)_ = 8.47, *P* < 0.01; *Fos*: *F*_(1,20)_ = 17.2, *P*=0.001, *Arc*: *F*_(1,20)_ = 8.72, *P* < 0.01] (***Figure 2*, *Figure 2 - Figure Supplement 4***). In contrast, repeated exposures to cocaine administration was characterized by an enhanced transcriptional induction in the LCtx, DS, and VTA. This enhancement was characterized by the significant induction of *Egr2* in the LCtx and DS and *Fos* in the LCtx, DS, VTA [interaction of treatment (cocaine vs saline) and time (acute vs repeated); *Egr2*: LCtx *F*_(1,29)_ = 6.43, *P* < 0.05; DS *F*_(1,29)=_ 4.58, *P* < 0.05; *Fos*: LCtx *F*_(1,29)_ = 5.35, *P* < 0.05; DS *F*_(1,29)=_ 4.21, *P* < 0.05, VTA *F*_(1,13)_ = 14.3, *P* < 0.01]. However, in the NAc, the initially robust induction of *Egr2* transcription following acute treatment decreased after repeated administration (interaction treatment x time, *Egr2*: F (1, 28) = 39.7, P<0.0001) (***Figure 2*, *Figure 2 - Figure Supplement 4***).

Repeated exposure to sugar was also represented by significantly enhanced transcription, most prominently in the LCtx [interaction of sucrose (sucrose vs water) and time (acute vs repeated); *Egr2: F*_(1,26)_ = 5.02, *P* < 0.05, *Fos: F*_(1,26)_ = 7.51, *P* = 0.01; *Arc: F*_(1,26)_ = 6.79, *P* < 0.05)] (***Figure 2*, *Figure 2 - Figure Supplement 4***). Furthermore, reinstatement of feeding was also represented by significant induction of IEGs in the LCtx, driven by transcription of *Egr2* and *Fos* (*Egr2: F*_(2,28)_ = 13.1, *P* < 0.0001; *Fos: F*_(2,31)_ = 41.5, *P* < 0.0001) (***Figure 2*, *Figure 2 - Figure Supplement 5, 6***). Interestingly, the experiences of repeated cocaine, repeated sucrose and reinstatement of feeding, though quite diverse in many affective and cognitive aspects, are all characterized by the reinforcement of an anticipated experience of positive valence. These results suggest that increased transcriptional representation, specifically in the LCtx, may serve as a correlate of growing salience of positively reinforcing experiences^14^. The transcriptional representation of reinforcement contrasts with the diminished transcriptional representation of anticipated and unavoidable aversive experiences^23^, providing correlates of positive reinforcement, in contrast to habituation.

### Decoding recent experiences of individual mice from minimal transcription

Finally, we tested if transcriptional signatures comprising a minimal set of gene markers could be used to decode the recent experience of individual mice. The transcriptional induction of five genes (*Arc, Egr2, Egr4, Fos and Fosb*) across five structures (LCtx, NAc, DS, Amy, LH) constitute 25 gene-structure ‘features’. Reasoning that not all 25 features are equivalent in the information they provide for decoding (defined as support), we performed Random K-Nearest Neighbor (R-KNN) feature selection^24^, to identify the features that provide maximal support (***Figure 3A***), and the minimal subset of features required to obtain the highest classification accuracy (***Figure 3B***). We identified that a combination of eight features (expression of *Egr2* and *Fos* in the LCtx, NAc and Amy, and expression of *Egr2* and *Fosb* in the DS) could enable the decoding of the recent experience of individual mice with an efficiency of 93.6% (***Figure 3C***). Random shuffling of the association of mice to experiences demonstrated the reliability of the classifier, and the potential for our results to generalize beyond the given dataset (***Figure 3D***). An intuitive representation of the differentiation of experiences based on particular features is provided by a decision tree (one of a number of possible trees), in which mice are manually assigned to appropriate branches according to the level of induction of a particular gene in a given structure (***Figure 3E***).

**Figure 3.**
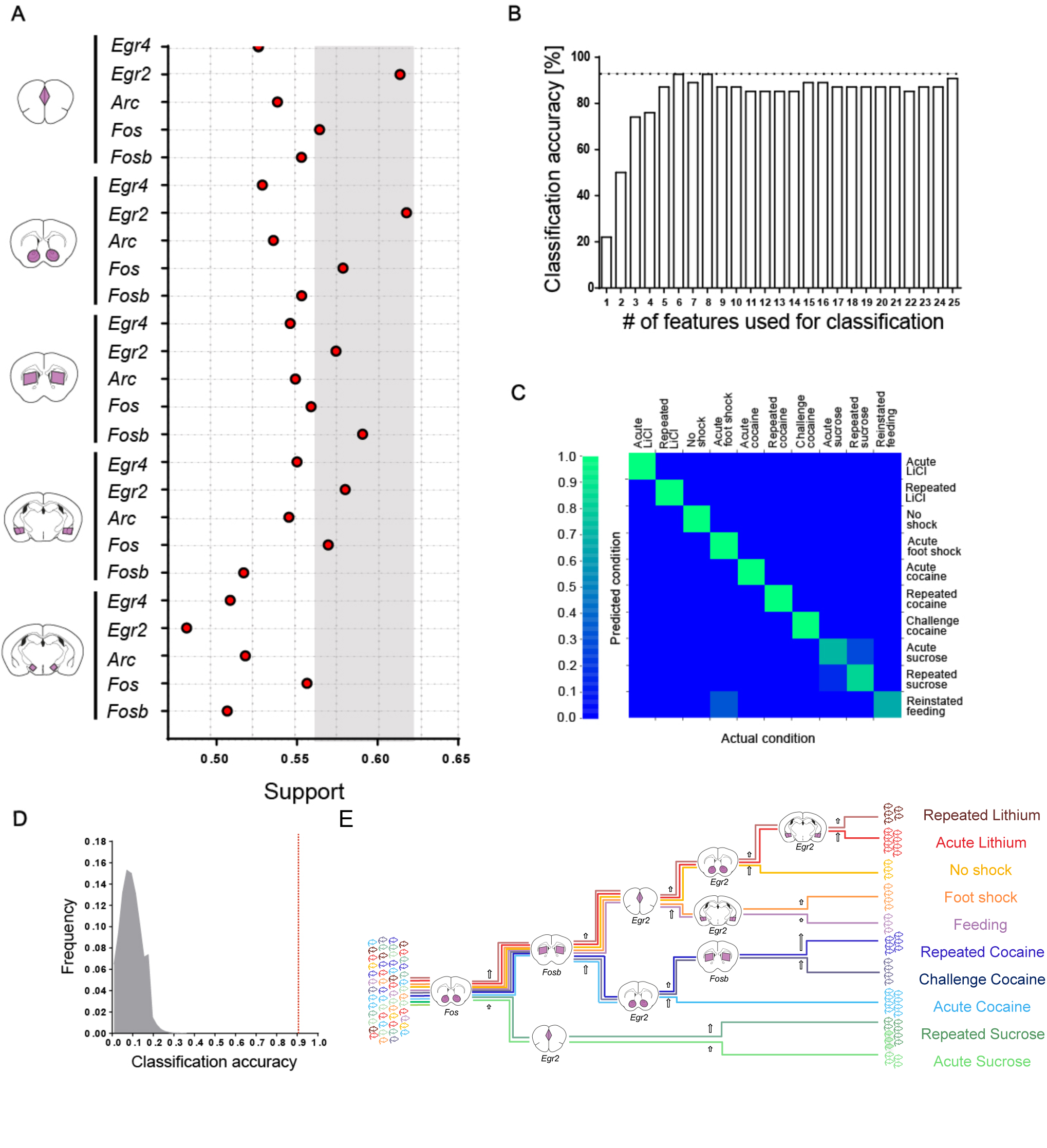
Decoding the recent experience of individual mice from minimal transcriptional signatures. **(A)** Selection of features (expression of IEG in a brain structure) with the highest potential to contribute to the classification of recent experiences (‘support’). Support was defined by running 1e^6^ classifiers with varying composition of features, and calculating the mean accuracy for each feature over all the iterations to which it contributed, using the Random k-Nearest Neighbors (RKNN) algorithm. The grey zone defines the eight features with the highest support values (*Egr2* and *Fos* in the LCtx, *Egr2* and *Fos* in the NAc, *Egr2* and *Fosb* in the DS, *Egr2* and *Fos* induction in the Amy). **(B)** Accuracy of classification obtained with an increase in the number of features. For each # (subset) of features, the features ranking highest in support were chosen and a KNN classifier was evaluated using only those features. Peak accuracy was achieved with eight features (93.6%). **(C)** Confusion matrix representing the classification accuracy (93.6%) of decoding the recent experience of individual mice based on the eight most informative features. Efficiency is scaled from blue to green, with bright green corresponding to 100% efficiency (n = 53 mice). **(D)** Verification of classification validity. A randomization test was performed, in which the classifier was run on 10^6^ random permutations of the association of individual mice to the appropriate experience, and the frequency of classification accuracies is plotted in grey, while the red dotted line represents the classification accuracy obtained for non-randomized data (93.6%). **(E)** A supervised decision tree enabling the classification of mice according to experience by minimal gene expression (many trees can equivalently segregate the data). Mice are classified manually based on features that enable segregation at each internal node. The decision tree failed in segregating three mice that experienced acute sucrose from the group of mice that experienced repeated sucrose. Mice are color-coded according to experience.

Taken together, these results establish that a minimal set of transcriptional markers form representative signatures of the recent experience of individual mice, and enable precise decoding of recent salient experiences at the resolution of individual mice.

## Discussion

The brain creates representations of the world, encoding salient information for long-term storage to support the development of adaptive behaviors. In real time, the representation of information has been shown to be correlated with neural activity in distinct brain structures^25^. Powerful demonstrations of the potential to decode sensory experiences and correlates of emotional state have been made in both rodents and humans from neural activation patterns using in-vivo electrophysiology, fMRI, and other imaging techniques^26–30^. In this study, we demonstrate that IEG expression data from multiple regions of the mouse brain enables the decoding of recent salient experiences with high precision. We report that beyond mere ‘activity markers’ for labelling neurons activated during an experience, IEG expression provides a quantitative and scalable metric, representing a neural transcriptional code of recent experience. Interestingly, this neural transcriptional code is comprised of distinct transcriptional signatures that correlate to the affective attributes of the experiences that are being encoded, such as salience and valence. Moreover, these IEG expression patterns are modulated following repeated administration of a stimulus of a given value (positive, negative or neutral), suggesting a role for inducible transcription in sustaining long-term adaptations underlying the development of adaptive behavior. As this code is comprised of molecular components, it also provides a rich resource for biological insight into the processes underlying the long-term encoding of experience-dependent plasticity.

Transcriptional markers have been successfully utilized for the classification of developmental stages^31^, diseases^32,33^, and many other aspects of contemporary biomedical science^34^. Here we describe the utility of transcriptional markers for classification of salient experiences characterized by diverse affective properties. While the information embedded in the expression pattern of a single gene is not sufficient, a minimal subset of transcriptional markers enables the decoding of recent experience with high accuracy. Importantly, the principles we identify likely generalize to a broad set of experiences. However, it is also likely that the markers we utilize in our study could be (at least in part) substituted by other markers, depending on the choice of classifier, providing comparable results.

According to the Russell circumplex model^11,12^, affect can be defined in two dimensions – valence and salience. Valence has been suggested to be encoded in the Amy, PFC, NAc and VTA^35^. Our results demonstrate that experiences of negative valence are represented by a distinct transcriptional induction in the Amy. In contrast, experiences with positive valence induce transcription in the LCtx, NAc, DS and VTA. Moreover, we report that upon repetition, the transcriptional representation within these structures is dynamically modulated, potentially underlying long term adaptations following positive and negative reinforcement. Taken together our results proposes that inducible transcription is a rich resource for the identification of the involvement of specific brain regions in encoding affective properties of an experience, providing biological insight into the molecular processes underlying experience-dependent plasticity.

To explain how changes in transcription could affect future behavior, we introduce the concept of ‘predictive transcriptional coding’. Predictive transcriptional coding frames inducible transcription not as a reporter of a recent event, but rather as encoding the valuation of the experience. This experience-dependent plasticity, mediated by transcription, sets the state of the network in the context of a particular experience, priming it for prospective network plasticity, and adjusts the future response of the individual to the occurrence of a similar event. This notion is conceptually similar to the ‘reward prediction error’^36^, but is established on prolonged time scales. In this context, transcription also serves as a ‘salience filter’ – defining whether an experience is significant enough to induce plasticity and be encoded for long-term storage. Our results are consistent with transcription serving as a salience filter, whereas the valuation of a recent experience is encoded by the identity of the neural circuits recruited by the experience and the magnitude of transcription induced within them. A crucial question arising from this concept is: how do neurons or neural networks determine the threshold to commit to induction of transcription? One possibility could be that the threshold for commitment to transcription depends on coincidence of glutamatergic and neuromodulatory inputs. It should be noted that our study revitalizes concepts and questions, which were raised previously in a landmark treatise defining inducible transcription as a ‘genomic action potential’ for encoding experience^37^.

Our work provides a numerical definition of the imprint of recent experience, demonstrating a quantitative and predictive approach for analysis of neuronal plasticity underlying adaptive behavior. Quantitative definitions of interoceptive states are expected to have implications for drug development - providing objective metrics for comprehensive characterization of the individual perception and valuation ascribed to an experience by individual subjects. For example, in the context of abuse liability, an objective quantitative interoceptive metric of the hedonic potential of a compound could increase standardization, reducing the reliance on variable behavioral outcomes.

While there is substantial investment being made in the development of methodologies for transcriptional profiling with deeper coverage and increasing spatial resolution, our study demonstrates that fundamental phenomena can be identified by applying simple methods with low spatial resolution and coverage. Future work, applying tools of higher resolution, could build on our observations to address additional questions – such as the spatial distribution of neuronal ensembles recruited by experience and the identity of cell types recruited by distinct experiences.

Approaches for non-invasive quantitative measurement of the encoding of experience can be envisioned, utilizing fluorescent markers of inducible transcription in combination with whole-brain imaging^38^. New technologies are rapidly emerging for whole-brain analyses of transcription^39–41^, as are strategies for comprehensive profiling of single neurons^42,43^. These technological developments, together with the novel concept we develop here, are expected to provide the foundation for a new area of neuroscience research. This discipline, of “Behavioral Transcriptomics”, will apply transcriptional analysis for investigation of intricate mechanisms of neural circuit plasticity underlying cognition. We propose that the approach of behavioral transcriptomics will provide a systems-level view of the encoding of experiences to long-term memory. Potentially, different elements of an experience may be mediated by activation of defined receptor sub-types (e.g. NMDA, dopamine, serotonin, etc.’) at specific dendritic locations, each inducing a component of the transcriptional program. If so, taken to its extreme, deciphering this transcriptional code will enable precise decoding of synapse-specific plasticity from quantitative analysis of inducible transcriptional markers.

## Materials and Methods

### Animals

Male C57BL/6 mice aged 6-8 weeks (Harlan Laboratories, Jerusalem, Israel) served as subjects for the study. Mouse body mass ranged from 18 to 35 g, while between experimental groups in each repetition of experiments, the difference in body mass between animals did not exceed 4 grams. Four to five mice were housed per cage in all experiments except for sucrose consumption experiments, for which animals were single-housed. Mice were maintained in 12-12 h light/dark cycle (0700 on/ 1900 off), in a temperature (20–22°C) and humidity (55 ± 10 %) controlled facility. Mice received ad libitum access to water and food, with the exception of the experiment studying reinstatement of feeding, in which they were food deprived for 18 hrs before reinstatement of feeding. Mice were randomly assigned to experimental groups and tested according to Latin square design. All tests were conducted during the light phase of the circadian cycle. Each experiment was performed at least twice, by independent researchers in the group, and provided similar results. All animal protocols were approved by the Institutional Animal Care and Use Committees at the Hebrew University of Jerusalem and were in accordance with the National Institutes of Health Guide for the Care and Use of Laboratory Animals. A table defining the number of mice (‘n’) contributing to each figure is included as Supplementary Table 2.

### Behavioral Assays

Mice were acclimated to the animal facility for at least 2-5 days, followed by 3-4 days of experimenter handling, before the start of an experiment. Maintenance of uniform conditions across experiments and extensive handling were essential for reducing experimental variability, enabling the identification of a robust transcriptional response specifically induced by the experience being tested and minimal contamination from contextual background.

### Behavioral sensitization to cocaine

Mice were subjected to three days of intraperitoneal (i.p.) saline injections (250 microliter/injection), prior to exposure to cocaine (20 mg/kg freshly dissolved in physiological saline to 2 mg/ml and injected at a volume of 10 ml/kg; cocaine was obtained from the pharmacy at Hadassah Hospital, Jerusalem). The *acute cocaine* group received a single i.p. dose of cocaine, followed by analysis of locomotor behavior for 15 minutes in a video-monitored open-field arena. Animals were finally taken from their home cage and sacrificed at 1, 2 and 4 hrs following the cocaine injection. The *repeated cocaine* group received five consecutive daily injections of cocaine, and were studied (similar to the acute cocaine group) following the fifth cocaine injection. The *challenge cocaine* group were treated as the repeated cocaine group, and then made abstinent from cocaine for 21-22 days, following which they were challenged with cocaine and re-exposed to the open-field arena. All responses were normalized to baseline controls (time 0), which were interleaved with their peer group, but were not treated on the day of the experiment. Additional reference groups included *acute saline without habituation,* which were habituated to the open-field arena for three days after a brief period of handling, and were sacrificed 1 hr following a single injection of saline. Responses in this group were normalized to controls (time 0), which were not exposed to any saline injections. The group of *acute saline without habituation* served as a reference for the habituation of the *acute saline* group, in which animals were treated identically to the acute cocaine group (i.e. three consecutive days of habituation to saline injections in the open-field arena), but received a saline injection on the day of the experiment. Following each i.p. injection, mice were placed in an open-field arena for 20 minutes, during which locomotion was assayed between minutes 2 to 17.

### LiCl exposure

All mice were habituated to injections of saline and locomotor monitoring in an open-field arena for three days preceding onset of the experiment. Animals were subjected to either acute or repeated administration of LiCl (Sigma-Aldrich, St.Louis, MO, USA). In *acute LiCl* experiments, mice were administered with either a single dose of LiCl (at 150 or 250 mg/kg, as indicated in figure legends) or saline. In the experiments testing *repeated LiCl*, mice received LiCl (150 mg/kg) for five consecutive days, and following a 48 hr break were re-exposed to LiCl or saline. Mice were divided into four groups: a) Received saline injections for five days and were not exposed to an injection on the last day (*saline-0h*), b) Received LiCl injections for five days and were not exposed to an injection on the last day (*LiCl-0h*), c) Received saline injections for five days and were subjected to saline injection on the last day (*repeated saline*), d) Received LiCl injections for five days and were exposed to LiCl injection on the last day (*repeated LiCl*). In all experiments, immediately following administration of LiCl or saline, mice were placed in video-monitored open-field arenas for 30 min.

### Reinstatement of feeding

Mice were food deprived for 18 hrs before the experiment and then re-exposed to food for 1, 2 or 4 hours before they were sacrificed. Control animals (0 hr) were sacrificed immediately after the 18 hr food restriction. An additional reference group was allowed to continuously feed.

### Sucrose Consumption

Mice were single-housed for at least seven days before the experiment and habituated to the addition of a second water bottle in the cage for three days before the onset of the experiment. *Acute exposure to sucrose* was tested by habituating mice to the bottle with 10% sucrose overnight (16 hr), and 48 hrs later, re-exposing the mice to a bottle with sucrose or water (control) for 1 hr. *Repeated exposure to sucrose* was tested by exposing mice to sucrose repeatedly for eight consecutive days, 2 hr each day (12:00-14:00), and after a 48 hr break, re-exposed to sucrose or water (control) for 1 hr. Mice were sacrificed 1 hr following the exposure to sucrose. Sucrose and water intake were measured as a test for sucrose preference over water.

### Foot Shock

Following habituation to the experimental setup, the mice were placed in the experimental chamber (20 x 18 cm) for three minutes, during which time, baseline freezing behavior was measured. At three minutes, each subject received three mild foot shocks (2 s, 0.7 mA) separated by 30 s interval and post-shock freezing behavior was assessed immediately thereafter for 30 seconds before return to the home cage. Freezing, defined as a lack of movement other than respiration, was measured using Ethovision software (Noldus, Wageningen, The Netherlands).

### Locomotor activity measurement

Locomotor activity was assessed in sound- and light-attenuated open-field chambers. Mice were placed individually in a clear, dimly lit Plexiglas box (30×30×30 cm) immediately after injection of cocaine, LiCl or saline. Activity was monitored with an overhead video camera for 20 or 30 min (in cocaine sensitization and LiCl experiments respectively) using Ethovision software (Noldus, Wageningen, The Netherlands).

### Dissections

Performed as previously described^44^. Mice were deeply anesthetized with Isoflurane (Piramal Critical Care, Bethlehem, PA, USA) and euthanized by cervical dislocation, followed by rapid decapitation and harvesting of brains into ice cold artificial cerebrospinal fluid (ACSF) solution (204 mM sucrose, 26 mM NaHCO3, 10 mM glucose, 2.5 mM KCl, 1 mM NaH_2_PO_4_, 4 mM MgSO_4_ and 1mM CaCl_2_; all from Sigma-Aldrich, St. Louis, MO). Coronal slices (400 µm) were cut on a vibrating microtome 7000 smz2 (Camden Instruments, Loughborough, UK) in ice-cold artificial cerebrospinal fluid (ACSF). Brain regions [Limbic cortex (LCtx), Nucleus Accumbens (NAc), Dorsal Striatum (DS), Amygdala (Amy), Lateral Hypothalamus (LH), Hippocampus (Hipp) and Ventral Tegmental Area (VTA)] were dissected from relevant slices under a stereoscope (Olympus, Shinjuku, Tokyo, Japan). Samples of LCtx, NAc, DS, Amy, LH AND Hipp were obtained from 2* 400 µm thick sections, while VTA, was obtained from 2* 200 µm thick sections (***Methods Section – Figure Supplement 1***). All of the steps were performed in strictly cold conditions (∼4°C) and care was taken to avoid warming of the tissue sections or the ASCF at all times. The tissue pieces were immediately submerged in Tri-Reagent (Sigma-Aldrich, St.Louis, MO) and stored at -80°C until processing for RNA extraction.

### Marker selection, RNA extraction, qPCR and microfluidic qPCR

The strategy for marker selection consisted of three steps. The initial list of candidate IEGs was compiled from a whole-genome microarray analysis of transcriptional dynamics induced by cocaine experiences in the nucleus accumbens (Illumina MouseRef-8 v2 Expression BeadChip microarrays; data not shown), as well as a survey of literature and databases pertaining to IEG expression. qPCR primer probes were developed for 212 genes and primer efficiency was tested, resulting in selection of 152 optimal primer pairs. Differential expression of the shortlisted IEGs was then tested on samples from multiple brain structures, dissected from mice following cocaine and LiCl experiences, utilizing microfluidic qPCR arrays. Genes that displayed at least 1.25-fold induction in any measurement were shortlisted, resulting in a list of 78 genes. Finally, the five genes that were induced most consistently across all structures and conditions were selected for analysis by traditional qPCR.

RNA extraction was performed strictly in cold RNase-free conditions. Tissue was homogenized using a 25G needle attached to a 1 ml syringe or using TissueLyser LT (Qiagen, Redwood city, CA, USA). The homogenate was centrifuged at high speed (15kg for 10 min) and the supernatant was mixed with chloroform (Bio-Lab, Jerusalem, Israel) by vigorous shaking and centrifuged (15k g for 15 min) to separate the RNA from other nucleic acids and proteins. Isopropanol (J. T. Baker, Center Valley, PA) and glycogen (Roche, Basel, Switzerland) were added to the aqueous layer and samples were placed either at -20°C for 24 hours or at -80°C for 1 hour (producing comparable results). The samples were centrifuged at high speed (15k g for fifteen min) for the precipitation of the RNA. The RNA was then washed in 75% ethanol (J. T. Baker, Center Valley, PA) by centrifugation (12k g for five min), dried and dissolved in ultrapure RNase free water (Biological Industries, Kibbutz Beit Haemek, Israel). RNA concentration was measured with a NanoDrop 2000c spectrophotometer (Thermo, Wilmington, DE) and random-primed cDNA was prepared from 100-300 ng of RNA, with use of a High Capacity cDNA Reverse Transcription Kit (Applied Biosystems, Foster city, CA), following manufacturer guidelines.

cDNA was processed for qPCR analysis using qPCR primer pairs (IDTDNA, Coralville, IA) and SYBR Green in a Light-cycler® 480 Real Time PCR Instrument (Roche Light Cycler*480 SYBR Green I Master, Roche, Basel, Switzerland) according to manufacturer guidelines. Relative levels of gene expression (ΔCt) were obtained by normalizing gene expression to a housekeeping gene (GAPDH). Fold induction was calculated using the ΔΔCt method, normalizing experimental groups to the average of a relevant control group.

Microfluidic qPCR, querying of 96 samples with limiting quantities of RNA against 96 sets of qPCR probes was performed utilizing Fluidigm Biomark Dynamic IFC (integrated fluidic circuit) Arrays (Fluidigm Corp, South San Francisco, CA). Briefly, samples are subjected to targeted preamplification to enrich for specific gene products, which were then assayed with dynamic array fluidic microchips. Sample preparation was performed according to previously published protocols^44^. Targeted pre-amplification (STA) was achieved by mixing samples with a set of diluted primer pairs in TaqMan PreAmp Mastermix (Applied Biosystems; Foster City, CA, USA) followed by 10 minutes of denaturation at 95°C and 14 cycles of amplification (cycles of 95°C for 15s and 60°C for 4 min). Primers were then eliminated by use of ExoI exonuclease (NEB; Ipswich, MA), placed in a thermal cycler at 37°C for 30 mins and then at 80°C for 15 mins. Samples were then loaded onto a primed dynamic array for qPCR in a specialized thermal cycler [Fluidigm Biomark; Thermal mixing: 70°C for 40 min, 60°C for 30 s, 95°C denaturation for 60 s, followed by 40 cycles of PCR (96°C for 5 s, 60°C for 20 s)]. For data analysis, a reference set of genes was identified, whose expression remained constant across all experimental conditions (*Dkk3, Tagln3, Gars, Scrn1, Rpl36al, Mcfd2, Psma7 and Hpcla4*). In order to reduce the potential for introduction of experimental error by normalization to a single gene, a ‘global-normalization’ Ct value was created for each sample from the average Ct values of the genes within the reference set. Fold induction was calculated using the ΔΔCt method, normalizing each gene in a sample to the global-normalization value (ΔCt), followed by normalization of the experimental groups to the average of their relevant control group.

### Data analysis

All data are presented as mean ± standard error (s.e.m). Data were analyzed using t-tests, one-way or two-way analysis of variance (ANOVA), as appropriate. Tukey test was used for post hoc analyses of significant one-way ANOVAs. Multiple comparisons following two-way ANOVA were conducted with Bonferroni post hoc comparison. Differences were considered significant at the level of p < 0.05. Statistical analysis was performed, and bar graphs and line graphs were created, with Prism 6.0 (GraphPad, San Diego, CA). Heat maps were created in MATLAB R2012a (Mathworks, Natick, MA). Radar plots were created in Origin 6.0 (Originlab, Northampton, MA).

### Computational analyses

The analysis was performed on data obtained from 53 mice, each of which experienced one of the experiences (acute, repeated or challenge cocaine, acute and repeated sucrose, reinstatement of feeding, acute and repeated LiCl and foot shock and no-shock controls exposed to the same environment). Each mouse was represented by a vector of twenty five features [corresponding to the induction of five genes (*Arc, Egr2, Egr4, Fos* and *Fosb*) across five structures [limbic cortex (LCtx), nucleus accumbens (NAc), dorsal striatum (DS), amygdala (Amy) and lateral hypothalamus (LH)]. Each gene-structure combination was defined as a “feature”.

### Linear projection

Principal component analysis was performed on the induction level of *five genes* (*Arc, Egr2, Egr4, Fos* and *Fosb*) in three structures (LCtx, DS and Amy), reducing the multi-dimensional data to three dimensions. The resultant vector where each animal is represented by the principal components (PCA 1, 2 and 3) were plotted on a 3D plot using a code written in MATLAB R2015b (MathWorks, Natick, MA). Each dot on the plot corresponds to an individual animal, belonging to a specific experience group. Groups were represented by different colors according to the color code presented in ***Figure 2***.

### Supervised Classification

The classifier used was k-Nearest Neighbors (KNN), with k=1 over the Euclidean space. This approach was selected based on the observation that the transcriptional response of mice within an experience group formed unique clusters. We evaluated the performance of our classification by a leave-one-out method. In this approach, we iterated over each sample in our training set and classification was performed given the rest of the training set. Visualization of the accuracy of classification was performed using a confusion matrix, which conveys both mean precision and mean recall of each condition classified. After applying feature selection, using Random k-Nearest Neighbors (RKNN) as described below, we performed classification using a limited set of eight features (***Figure 3C)***. Codes were written in MATLAB R2015b (MathWorks, Natick, MA) and confusion matrices were created in Python using the Matplotlib library (http://matplotlib.org).

### Feature selection

Feature selections were performed using Random k-Nearest Neighbors (RKNN) algorithm^24^. The contribution of each feature for classification of individual experiences was called *support*. We chose large (n=1e^6^), random subsets out of the twenty five available features in varying sizes (between 1 and twenty five). For each such subset we trained a classifier. Each feature *f* appeared in some KNN classifiers, for example, set **C**(*f*) of size *M*, where *M* is the multiplicity of *f*. In turn, each classifier *c* ∈ **C**(*f*) is an evaluator of its *m* features. We defined the *support* of a feature *f* as the mean accuracy of all the classifiers in C(f). Namely:

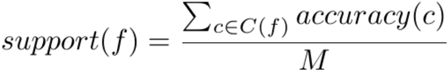

To further examine the effect of feature set sizes on classification performance we evaluated the classification accuracy of different subset sizes in the following manner: for each case, we chose the n features which were ranked the highest in their support, and evaluated the KNN classifier trained with those features only.

### Randomization

Considering the limited size of our dataset, we ensured that the classifier was not overfitted to our training set S. For this purpose, we produced a large number N (N=1e5) of permuted versions of our training set (s_i_,…s_N_), and created KNN classifiers in the same way we did for the original data. The permutation was performed by shuffling the association of individual mice with experiences. For each such permuted training set we learned a classifier and evaluated its classification accuracy (leave-one-out, see previous description). We calculated the empirical p value (p<1e-4) for the classification accuracy on our original training set in the following manner:

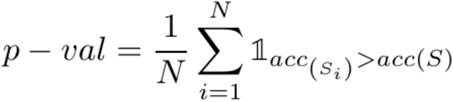

### Data and code availability

The data sets generated during the current study, as well as the code used for analysis are available from the corresponding author upon request.

---

**Figure 1 – Figure Supplement 1.**
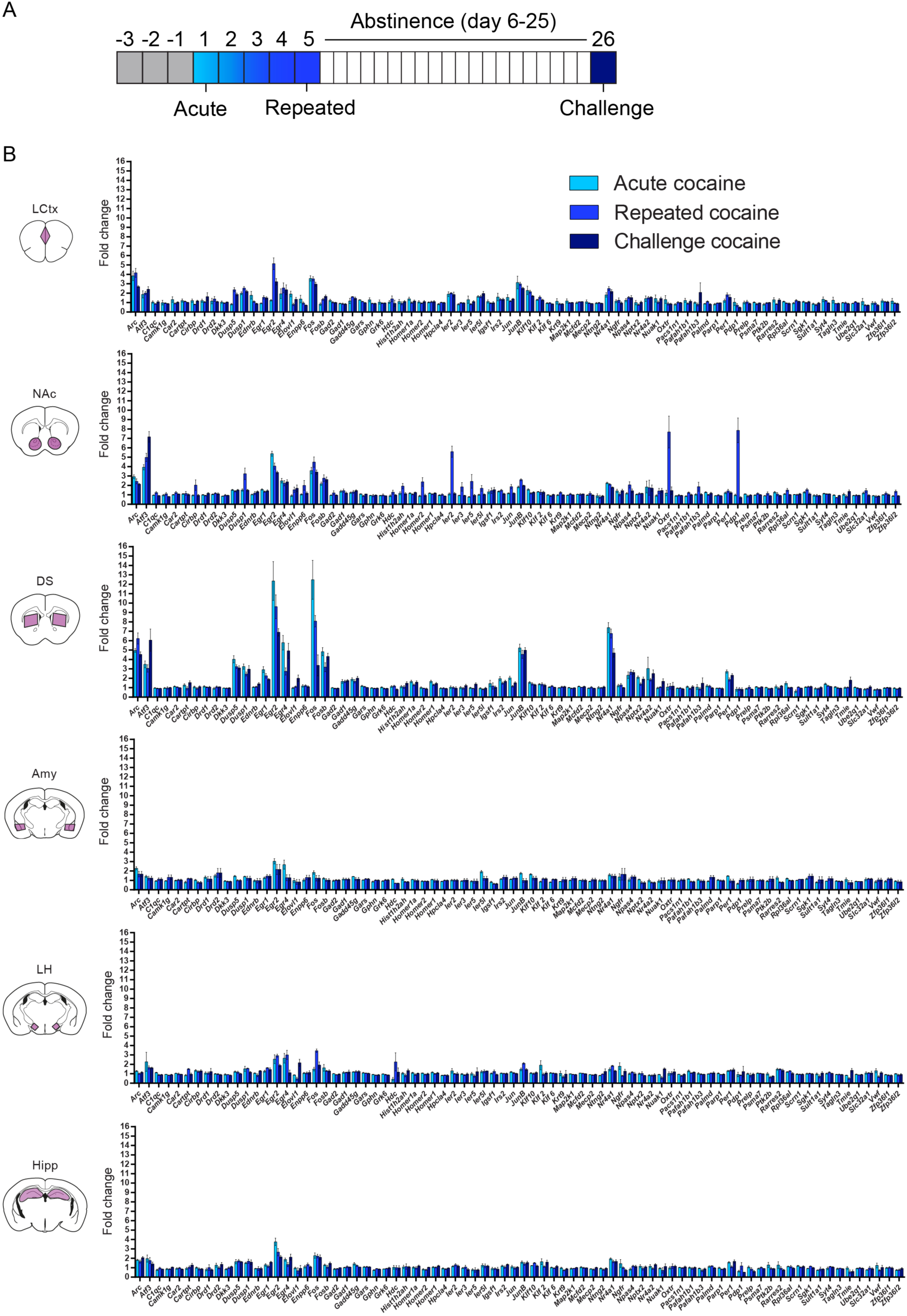
Acute, repeated and challenge cocaine experiences induce distinct transcriptional programs across brain structures. **(A)** Schematic of the experimental paradigm. **(B)** Average transcriptional induction of 78 genes one hour following acute, repeated or challenge cocaine experiences in 6 brain nuclei [LCtx: limbic cortex (n = 6-10), NAc: nucleus accumbens (n = 6-10), DS: dorsal striatum (n = 6-10), Amy: amygdala (n = 4), LH: lateral hypothalamus (n = 4), Hipp: hippocampus (n = 4)]. Genes are sorted alphabetically. Results indicate mean ± s.e.m.

**Figure 1 – Figure Supplement 2.**
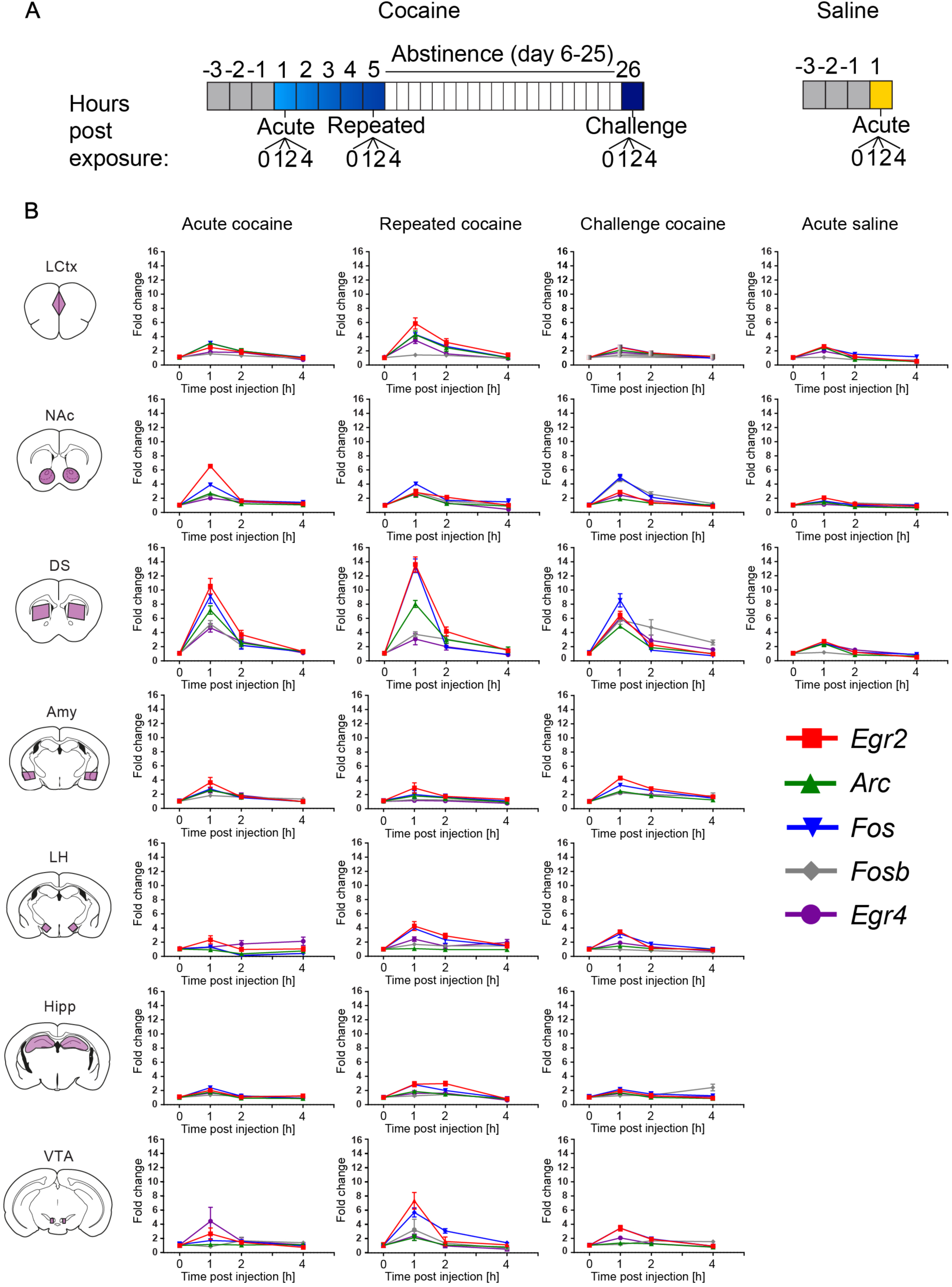
Time course of transcriptional induction of a minimal set of markers representing cocaine-induced transcriptional dynamics. **(A)** Schematic of the experimental paradigms. **(B)** *Arc*, *Egr2, Egr4, Fos* and *Fosb* induction (0, 1, 2, 4 hrs) following acute, repeated and challenge cocaine in seven brain nuclei (LCtx: limbic cortex, NAc: nucleus accumbens, DS: dorsal striatum, Amy: amygdala, LH: lateral hypothalamus, Hipp: hippocampus and VTA: ventral tegmental area), as well as acute saline (in the LCtx, NAc and DS). Subject numbers for each time point: n = 6-16 (LCtx, NAc and DS), n = 4-8 (Amy, LH and Hipp) and n = 2-4 (VTA) for acute, chronic and challenge cocaine. Acute saline included 6-8 animals in each time point for LCtx, NAc and DS. Results indicate mean ± s.e.m.

**Figure 1 – Figure Supplement 3.**
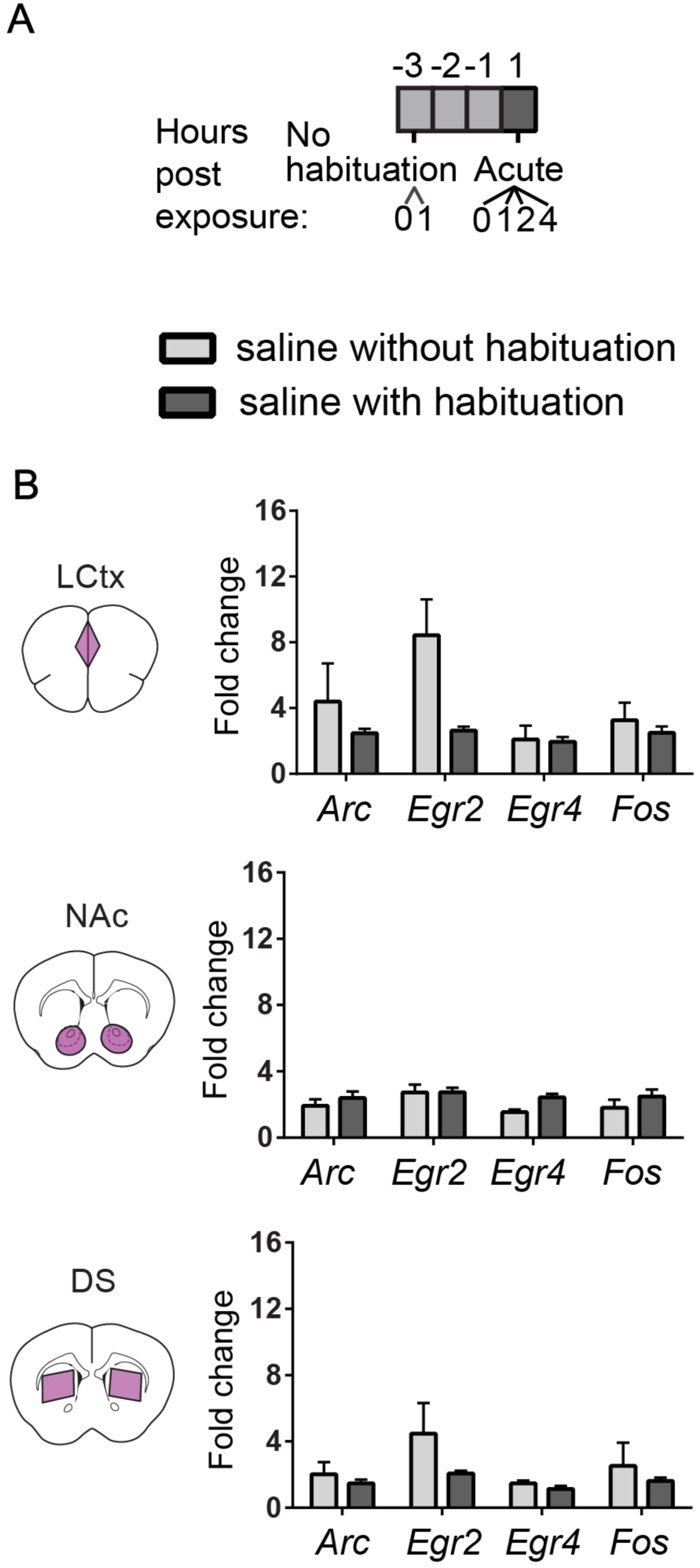
Characterization of repeated saline exposures illustrates the effect of habituation on induced transcription. **(A)** Experimental paradigm for i.p. saline injections. **(B)** Bar graphs demonstrating the transcriptional induction of *Arc*, *Egr2*, *Egr4* and *Fos* in the LCtx, NAc, and DS (limbic cortex, nucleus accumbens, and dorsal striatum, respectively) 1h following i.p. injection of saline with no prior habituation (n = 4), or following three days of habituation (n = 5). Results show mean ± s.e.m (in log_2_ scale).

**Figure 1 – Figure Supplement 4.**
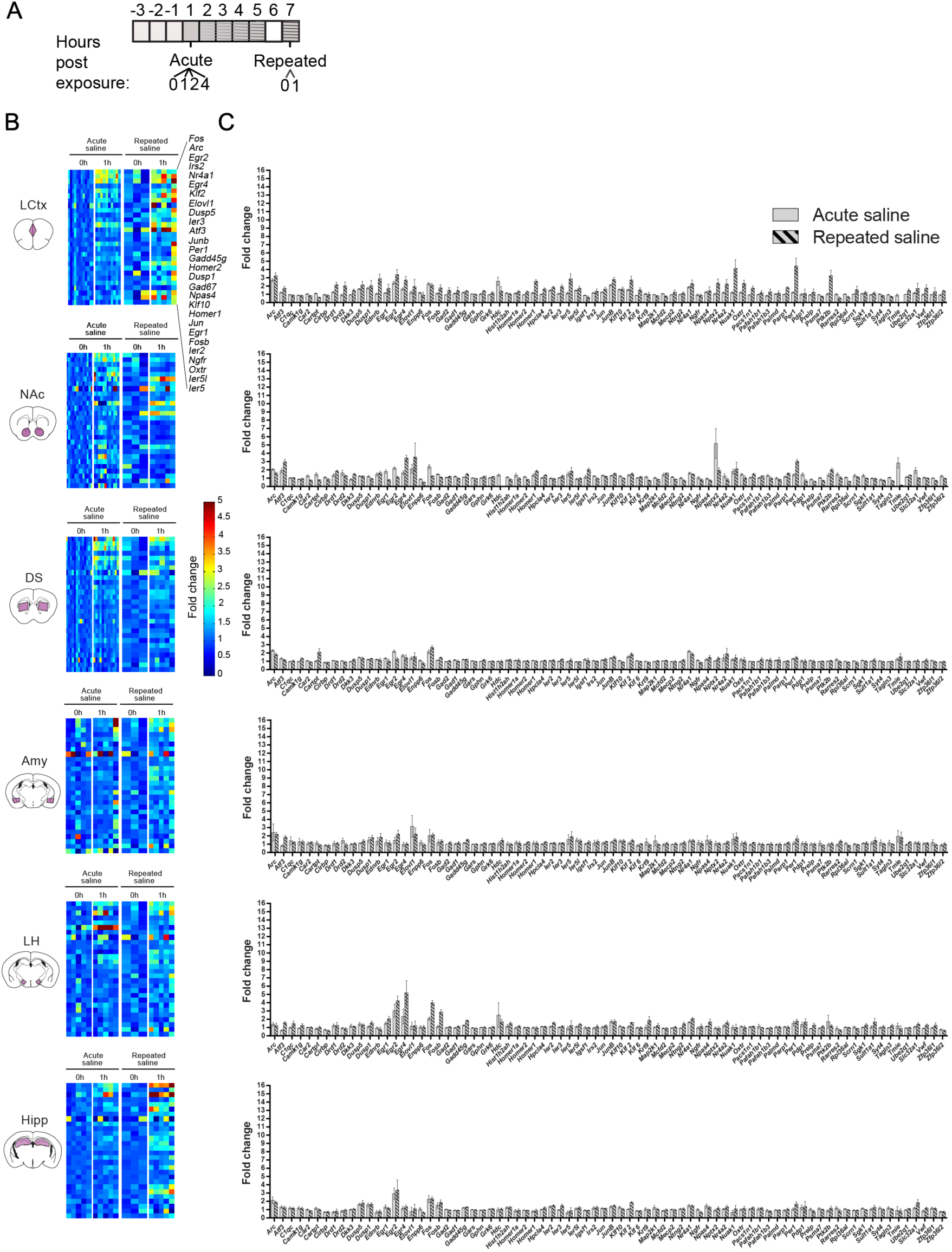
Characterization of the transcription induced by acute and repeated saline experiences. **(A)** Schematic of the experimental paradigm of i.p. injections of saline. **(B)** Expression matrix of the transcriptional induction of IEGs following acute or repeated saline experiences. Each column represents the transcriptional profile of an individual mouse. Sample numbers for each time point for acute saline: n = 10-12 in LCtx (limbic cortex); n = 12-13 in NAc (nucleus accumbens) and DS (dorsal striatum); n = 5 in Amy (amygdala), LH (lateral hypothalamus), Hipp (hippocampus). Sample numbers for each time point for repeated saline: n = 3-5 in LCtx, NAc, DS, Amy, LH and Hipp. Transcriptional induction is graded from blue (low) to red (high). Genes represented were induced on average at least 1.5-fold following either cocaine or LiCl experiences in any one of the studied brain nuclei. **(C)** Average transcriptional induction of 78 genes 1h following acute or repeated saline experience in the LCtx, NAc, DS, Amy, LH and Hipp. Genes are sorted alphabetically. Sample sizes: LCtx: n = 10 for acute saline and n = 5 for repeated saline; NAc: n = 12 for acute saline and n = 5 for repeated saline; DS: n = 12 for acute saline and n = 5 for repeated saline; Amy: n = 5 for acute saline and n = 5 for repeated saline; LH: n = 5 for acute saline and n = 5 for repeated saline; Hipp: n = 6 for acute saline and n = 5 for repeated saline. Results indicate mean ± s.e.m.

**Figure 2 – Figure Supplement 1.**
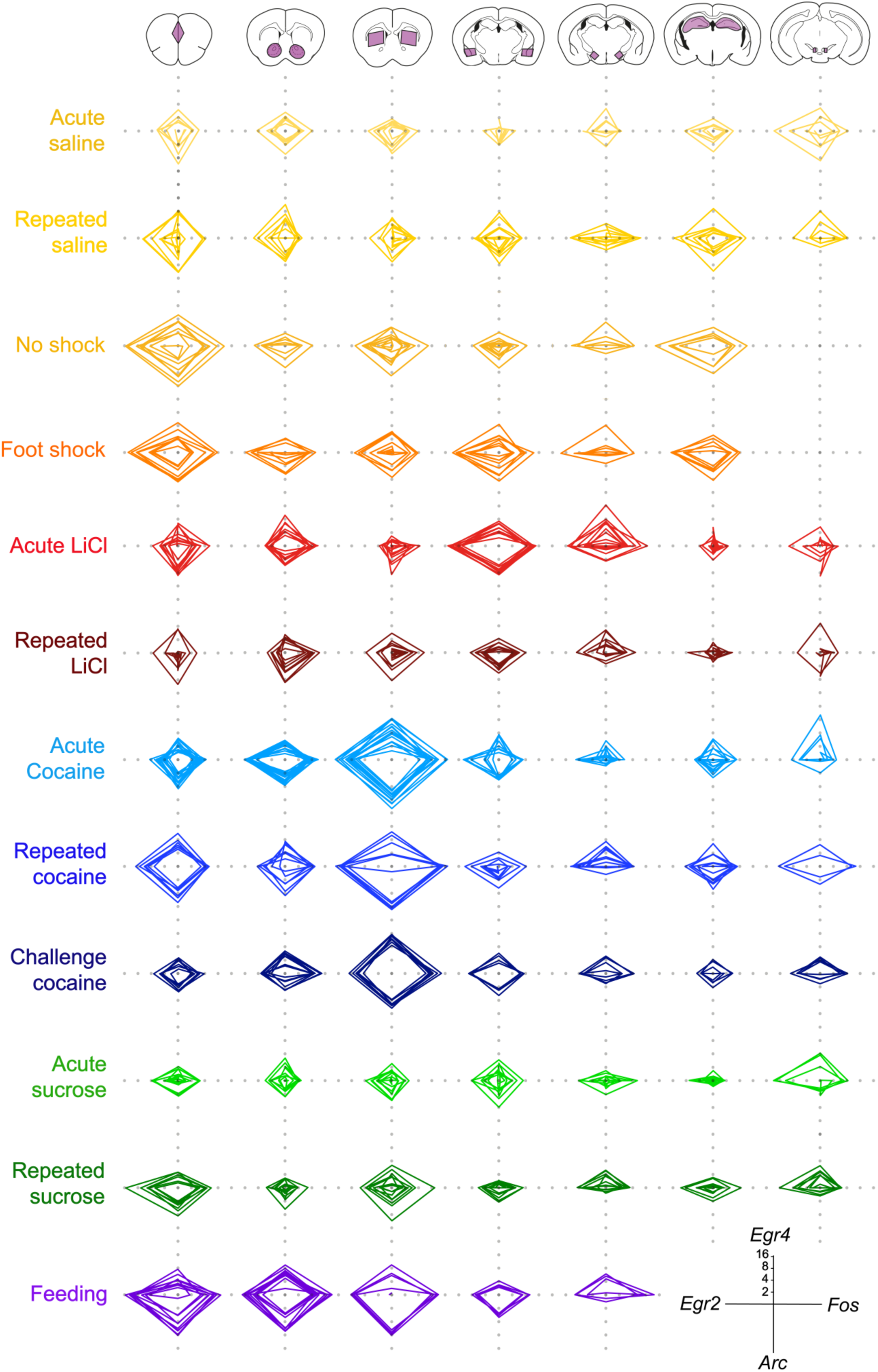
Low variance of the individual transcriptional representations of recent experience. Radar plots representing the transcriptional induction of *Arc*, *Egr2*, *Egr4* and *Fos*, in 7 brain nuclei [LCtx: limbic cortex (n = 4-14), NAc: nucleus accumbens (n = 4-14), DS: dorsal striatum (n = 4-14), Amy: amygdala (n = 4-9), LH: lateral hypothalamus (n = 3-9), Hipp: hippocampus (n = 4-8), VTA: ventral tegmental area, (n = 2-8)] 1 hour following the defined experiences. Results indicate fold induction at 1h in log_2_ scale, of individual subjects over relevant control conditions.

**Figure 2 – Figure Supplement 2.**
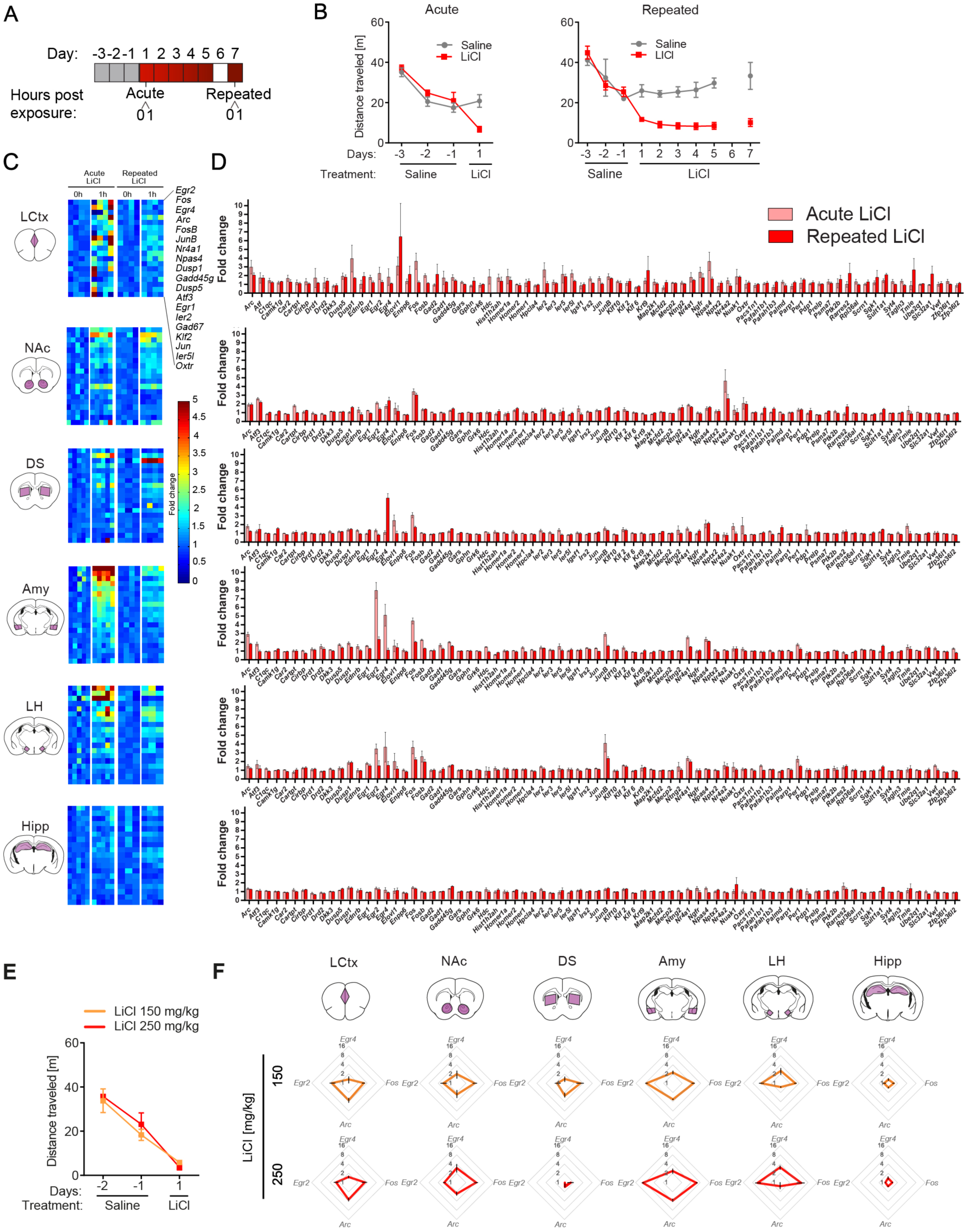
Acute aversive experiences are represented by robust induction of transcription in the amygdala. **(A)** Schematic of experimental paradigm. **(B)** Locomotor activity of mice following acute or repeated LiCl experiences, in comparison to saline. Sample size: acute saline, n = 6; acute LiCl, n = 9; repeated saline, n = 7; repeated LiCl, n = 8 mice per group. Results indicate mean ± s.e.m**. (C)** Expression matrix of the transcriptional induction of IEGs following acute or repeated experience of LiCl. Each column represents the gene induction profile of an individual mouse. Subject numbers for each time point following acute LiCl: n = 4 in LCtx (limbic cortex) and n = 5 in NAc (nucleus accumbens), DS (dorsal striatum), Amy (amygdala), LH (lateral hypothalamus) and Hipp (hippocampus). Following repeated LiCl: n = 4 in LCtx, NAc and Amy and n = 3-4 in DS, LH and Hipp. Transcriptional induction is graded from blue (low) to red (high). Genes represented were induced on average at least 1.5-fold over control in any one the studied brain areas. **(D)** Comparison of average transcriptional induction of 78 genes following acute LiCl (n = 4-5) or repeated LiCl (n = 3-4) at 1 hr following the experience, in the LCtx, NAc, DS, Amy, LH and Hipp. Genes are sorted alphabetically. Results indicate mean ± s.e.m. **(E)** Locomotor activity of mice following i.p. exposure to 150 mg/kg (n = 4) or 250 mg/kg LiCl (n = 4). **(F)** Radar plots representing the transcriptional response 1h following 150 mg/kg and 250 mg/kg of LiCl in the LCtx, NAc, DS, Amy, LH and Hipp. Results indicate mean ± s.e.m.

**Figure 2 – Figure Supplement 3.**
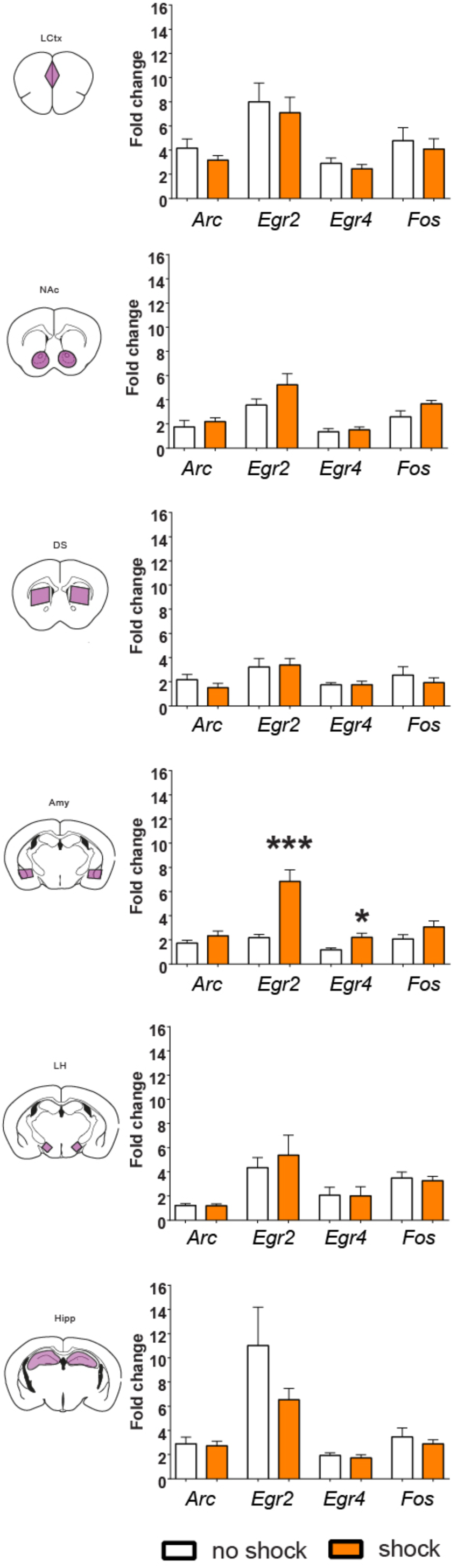
Transcriptional representation of negative valence in the amygdala. Comparison of average transcriptional induction of *Arc, Egr2, Egr4* and *Fos* in the LCtx, NAc, DS, Amy, LH and Hipp, 1 hr following acute exposure to the fear-conditioning chamber (no shock) or to a brief foot shock within the chamber (shock) Sample sizes across structures n = 3-8. Results indicate mean ± s.e.m. Student T-test: *p< 0.05, ***p<0.001 vs no shock control.

**Figure 2 – Figure Supplement 4.**
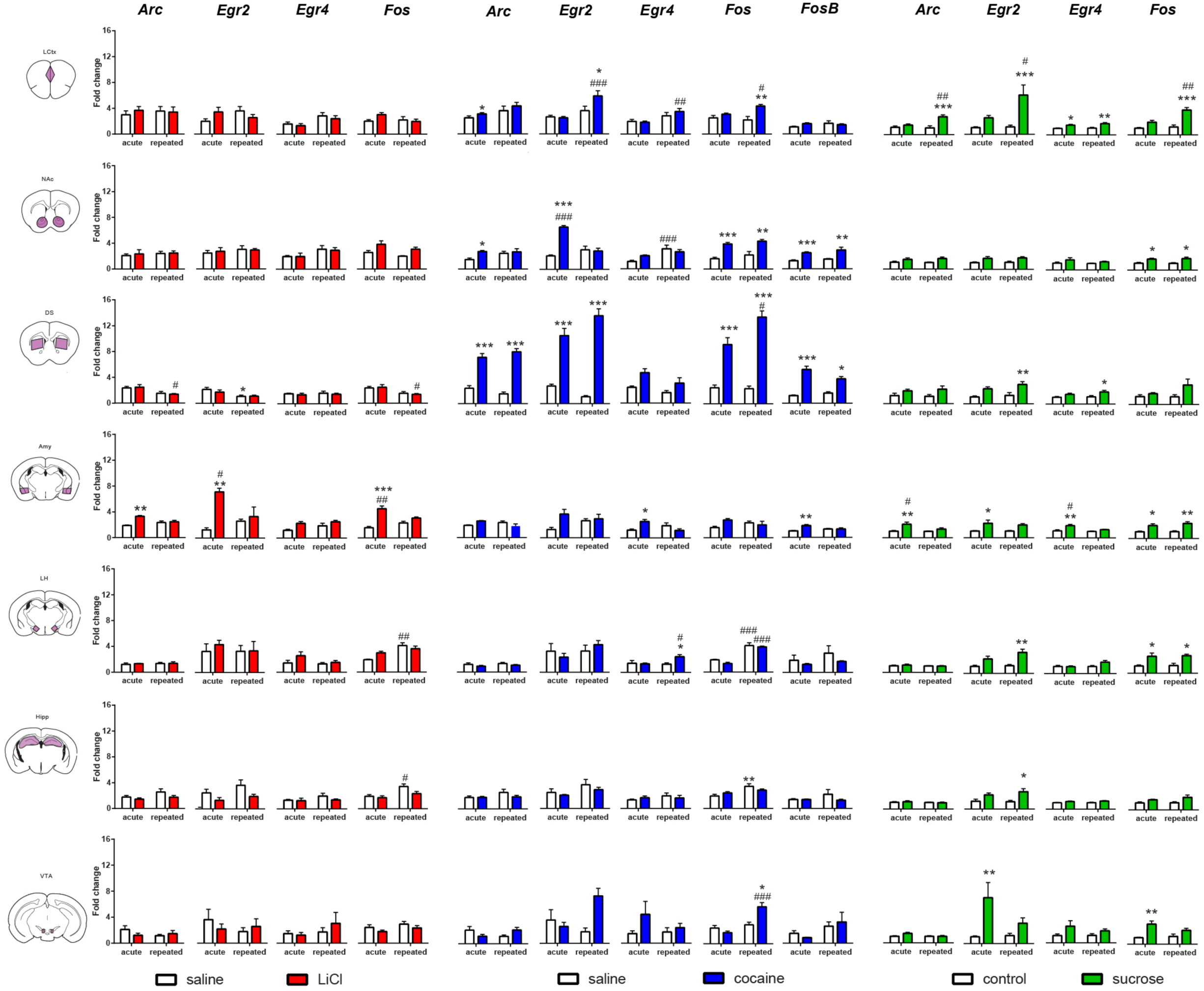
Transcriptional representation of habituation and reinforcement. Comparison of the transcriptional profiles induced by acute and repeated aversive (LiCl) and rewarding (cocaine and sucrose) experiences reveal distinctions and commonalities in the encoding of an anticipated experience of defined valence. Results indicate mean ± s.e.m Sample numbers for each time point-LCtx: limbic cortex (n = 6-14), NAc: nucleus accumbens (n = 6-14), DS: dorsal striatum (n = 6-14), Amy: amygdala (n = 4-9), LH: lateral hypothalamus (n = 3-9), Hipp: hippocampus (n = 4-9); VTA: ventral tegmental area (n = 2-8)]. Two-way ANOVA followed by Bonferroni post hoc comparison: *p<0.05, **p<0.01, ***p<0.001 vs control (saline or water), #p<0.05, #p<0.01, ###p<0.001 vs acute/repeated treatment.

**Figure 2 – Figure Supplement 5.**
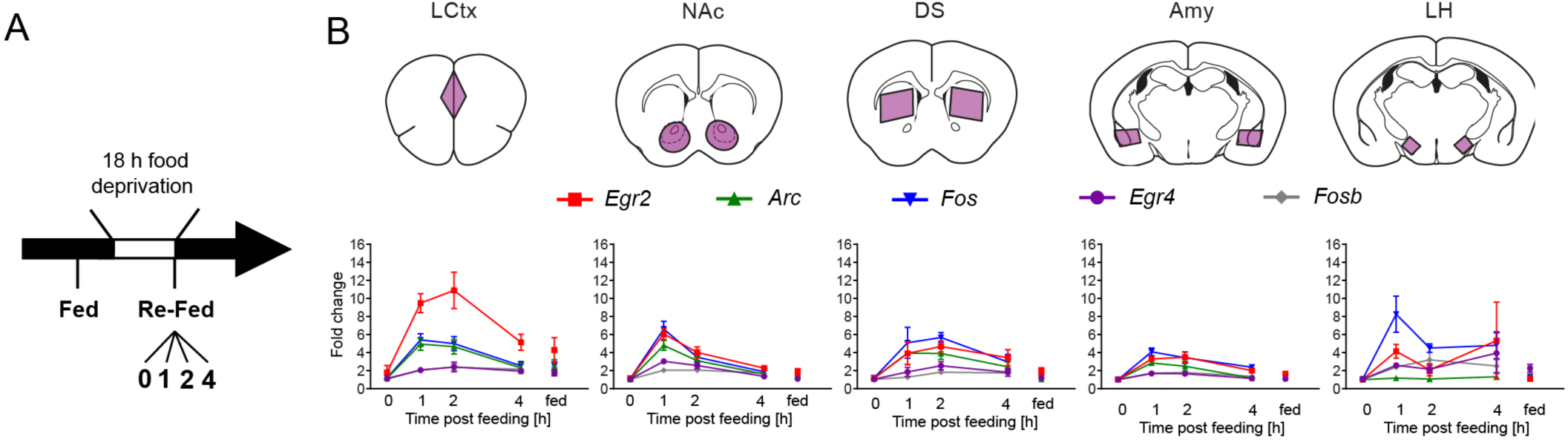
Reinstatement of feeding is represented by robust transcriptional dynamics. **(A)** Schematic of experimental paradigm for reinstatement of feeding. Mice were continuously fed or food deprived for 18 hrs, followed by analysis of transcription at 0, 1, 2, 4 hrs after reinstatement of feeding. **(B)** Time course (at 0, 1, 2, 4 hrs following reinstatement of feeding, in comparison to continuously fed mice, “fed”) of transcriptional induction of *Arc*, *Egr2, Egr4, Fos* and *Fosb* in 5 brain nuclei. Sample numbers for each time point: LCtx, limbic cortex n = 10-11; NAc, nucleus accumbens n = 10-11; DS, dorsal striatum n = 7-9; Amy, amygdala n = 3-4; LH, lateral hypothalamus n = 2-4. Results indicate mean ± s.e.m.

**Figure 2 – Figure Supplement 6.**
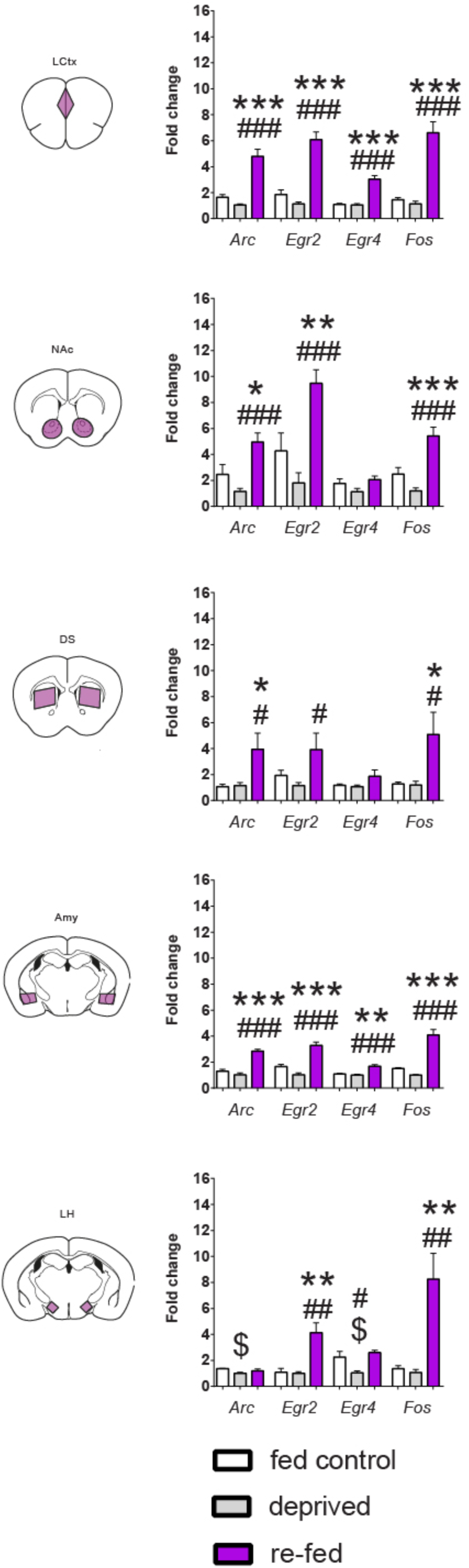
Reinstatement of feeding is represented by robust transcriptional dynamics. Comparison of transcriptional induction of *Arc*, *Egr2, Egr4 and Fos,* at 0 (deprived) and 1h following reinstatement of feeding (re-fed), and continuously fed mice (fed control) across 5 brain nuclei. Sample numbers for each time point: LCtx, limbic cortex n = 10-11; NAc, nucleus accumbens n = 10-11; DS, dorsal striatum n = 7-9; Amy, amygdala n = 3-4; LH, lateral hypothalamus n = 2-4. Results indicate mean ± s.e.m. One-way ANOVA followed by Tukey post hoc comparison: *p<0.05, **p<0.01, ***p<0.001 vs fed control; #p<0.05, ##p<0.01, ###p<0.001 vs deprived; $p<0.05 vs fed control.

**Methods Section – Figure Supplement 1.**
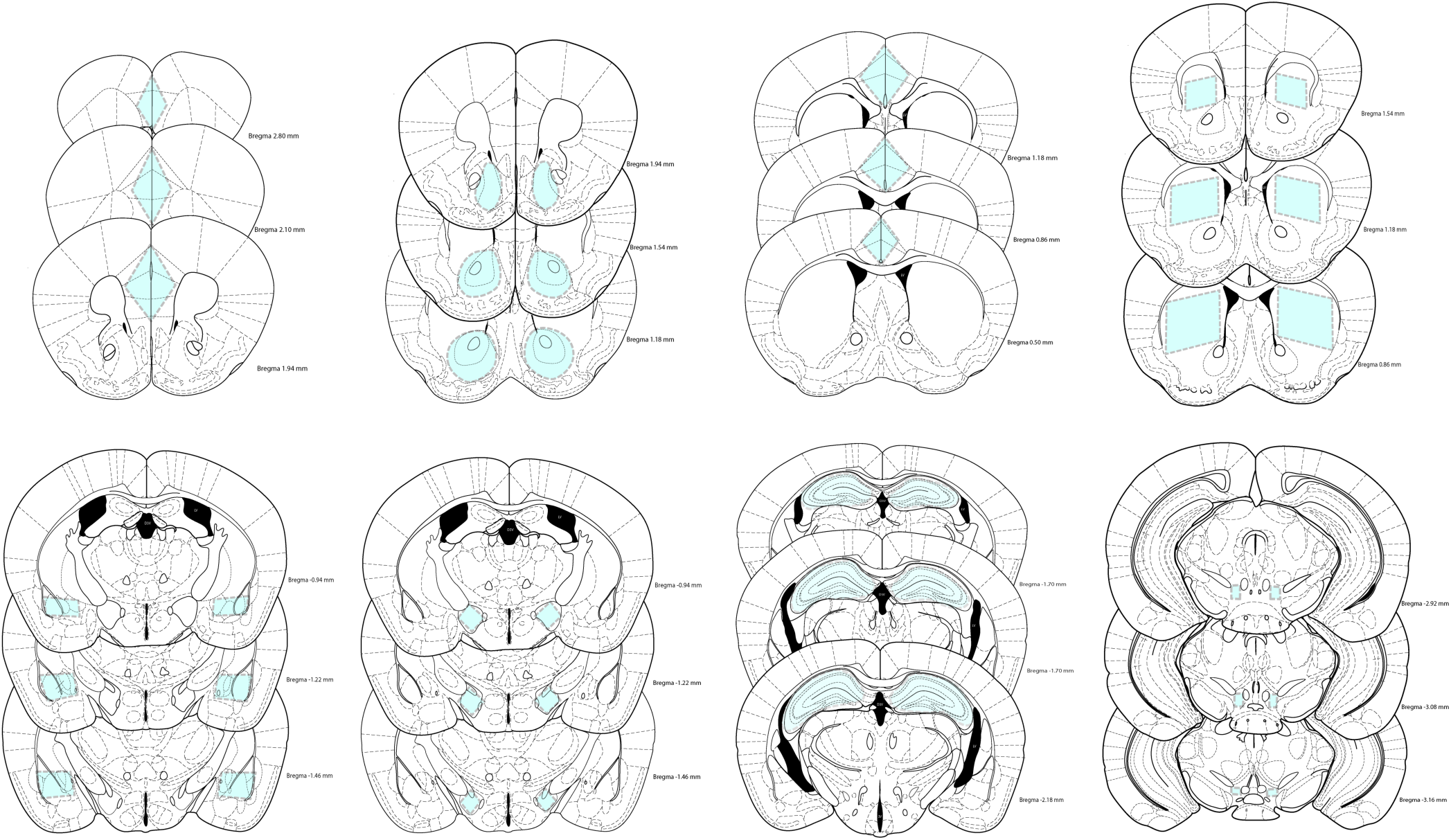
Boundaries of dissected structures. Shaded regions in blue represent areas cut out from 400 micron slices for LCtx (limbic cortex), NAc (nucleus accumbens), DS (dorsal striatum), Amy (amygdala), LH (lateral hypothalamus), and Hipp (hippocampus) and 200 micron slices for VTA (ventral tegmental area), providing reproducible tissue samples for analysis of transcription.

**Table S1**: Primer sequences and efficiency calculations.

**Table S2**: Animal numbers.

## Acknowledgements

This work was funded by grants to AC from the Israel Science Foundation (393/12 & 2341/15), ISF Center of Research Excellence on “Chromatin and RNA in Gene Regulation” (1796/12), EU Marie Curie (PCIG13-GA-2013-618201), the Brain and Behavior Foundation (NARSAD 18795), the German-Israel Foundation (2299-2291.1/2011), the Binational Israel-USA Foundation (2011266), the Milton Rosenbaum Endowment Fund for Research in Psychiatry, the Canadian Institute for Advanced Research, and contributions from Mr. Jaime Cohen (Mexico City) and the Stewart Resnick Foundation (Los Angeles). BIJ is funded by the Shimon Peres Fellowship from the Edmond and Lily Safra Center for Brain Sciences and the Lady Davis Foundation. Rob Malenka’s generosity in enabling preliminary studies to be performed in his laboratory is highly appreciated. We thank Hermona Soreq, Inbal Goshen, Mickey London, Sagiv Shifman, Zhiping Pang and members of the Citri lab for constructive criticism of the manuscript.

---

**Supplementary Table 1.**
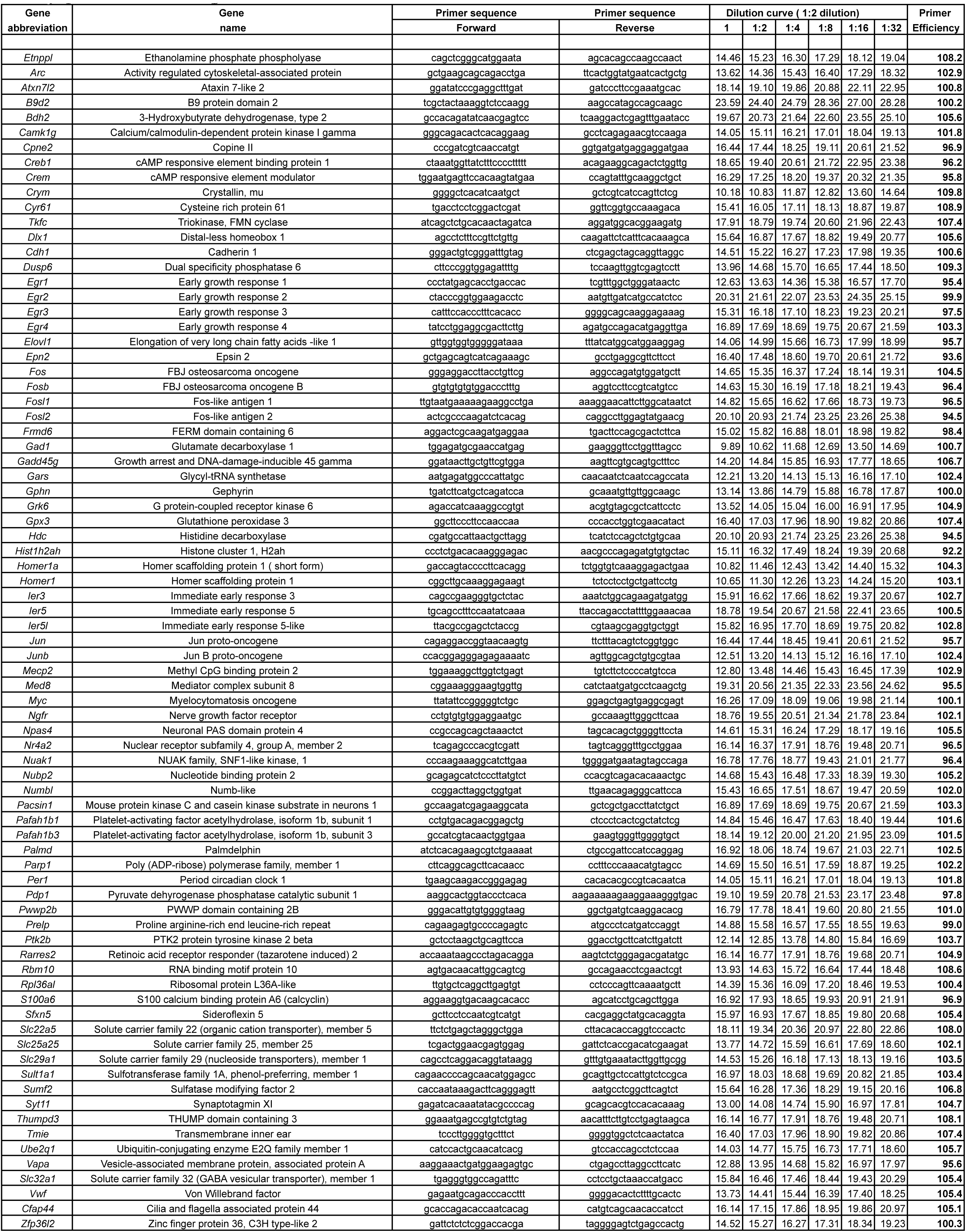
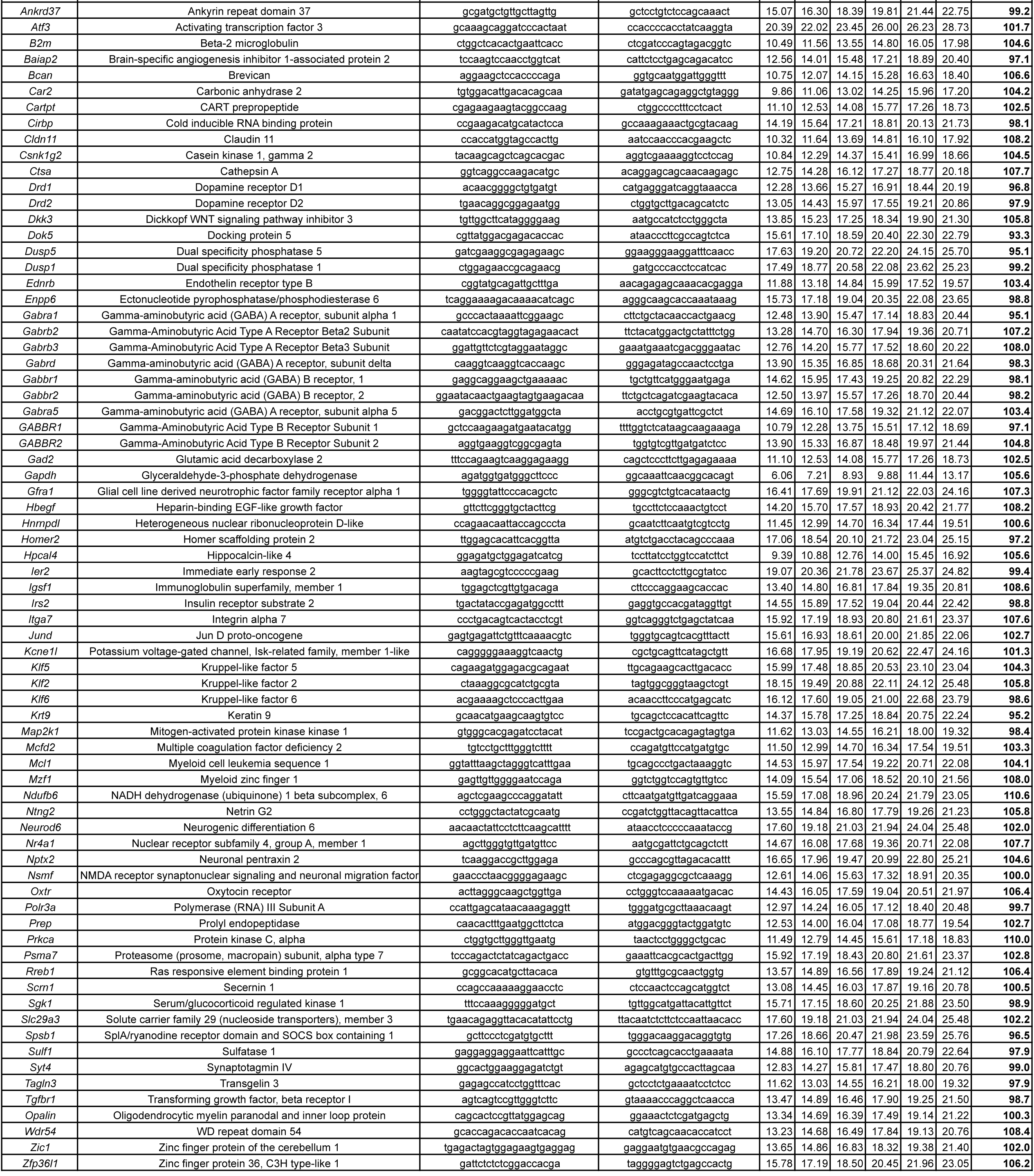
Primer characterization.

**Supplementary Table 2.**
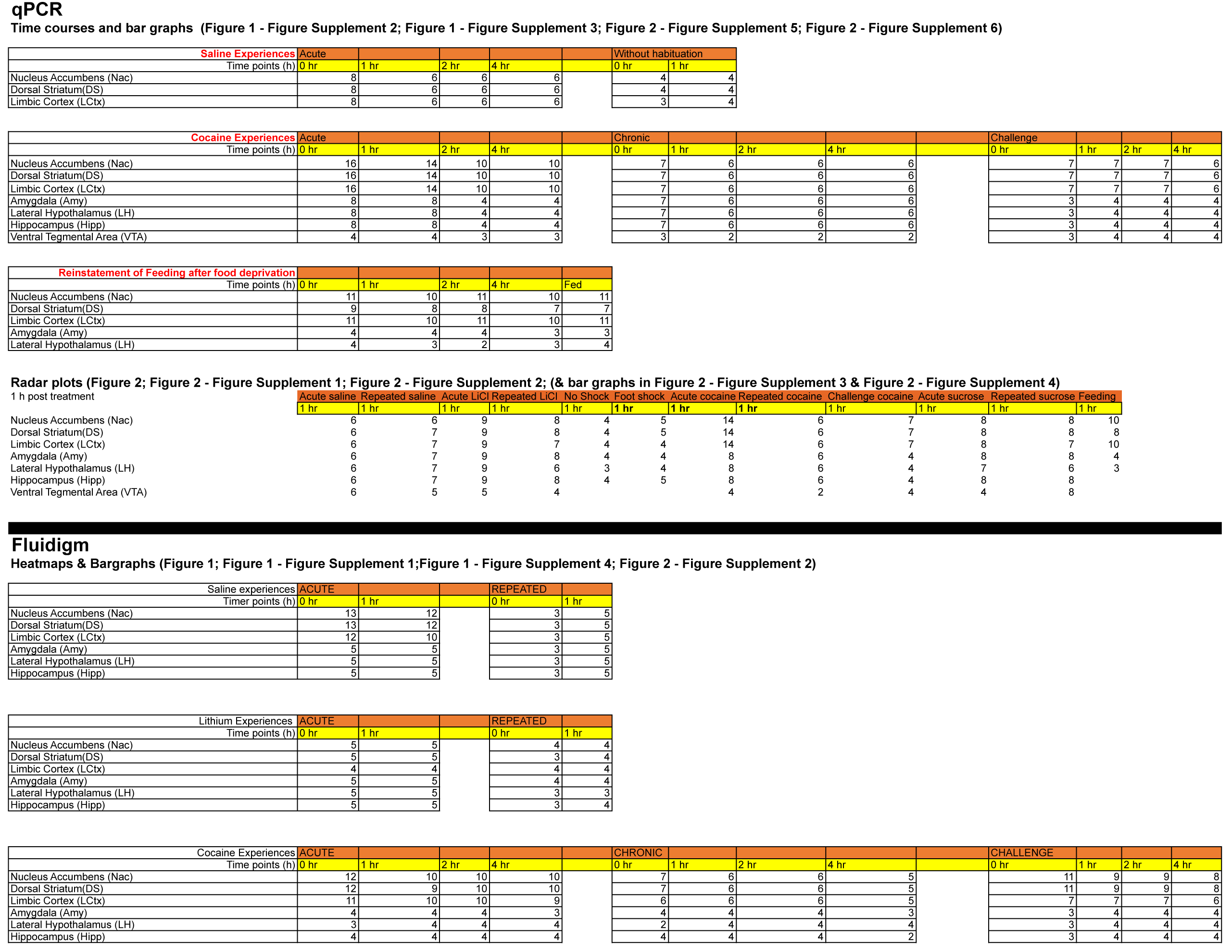
Animal numbers ('n')

